# Histone H2BK108Me2 Tunes Gluconeogenic Load in Type 2 Diabetes: A Molecular Dynamics Study

**DOI:** 10.64898/2026.01.12.699059

**Authors:** Ananya Chandra, Kharerin Hungyo

## Abstract

Type 2 Diabetes (T2D) is a serious metabolic disorder characterised by hyperglycemia, hyperinsulinemia, and insulin resistance. An increased rate of hepatic gluconeogenesis acts as one of the major contributors of the high blood glucose levels in the diseased condition. Transcriptional regulation is a main factor that controls gene expression. In this study we have investigated how the transcriptional availability of the *Cebpa* gene can be modulated at the nucleosomal level through the post-translational modifications (PTMs) of histones using a series of coarse-grained multi-microsecond molecular dynamics (MD) simulations. Our work explores the structural modulations imposed by histone PTMs on the *Cebpa* +1 nucleosome in terms of histone-DNA interactions, nucleosome unwrapping, and nucleosome sliding, which can further contribute to the alteration in transcription of the gene. The MD simulations reveal that the histone mark H2BK108Me2—a downregulated histone PTM mark found in a diet-induced obese mouse liver—can steer the nucleosome sliding in a direction such that the transcription start site of *Cebpa* gene tends to close. This study predicts H2BK108Me2 to be a potential histone PTM mark which might be involved in tackling gluconeogenic load by closure of nucleosomal DNA ends of *Cebpa* +1 nucleosome via sliding and posing chromatin unavailability towards essential transcription factors of the gene.

**Author Summary:** Histone PTMs regulate transcription of genes by altering nucleosome dynamics, yet, their precise mechanisms and effects remain unclear. Here, microsecond time-scale MD simulations with SIRAH forcefield reveals how T2D associated PTMs change DNA accessibility and reshape nucleosome conformation on the +1 nucleosome of a gluconeogenic regulator gene *Cebpa*. Our analysis uncovers changes in histone-DNA interactions, DNA trajectories, and nucleosome sliding to be the probable mechanisms of altered transcriptional output. This work attempts to bridge the gap between structural effects of nucleosomes and disease biology by providing a mechanistic link between metabolic disease epigenetics and chromatin biophysics. It demonstrates the role of PTMs in modulating the gene expression through collective nucleosome motions and offering insights into therapeutic targeting of histone PTMs.

## Introduction

Gluconeogenesis, the production of glucose by the liver from non-carbohydrate substrates, is one of the most important factors in causing the constantly elevated blood glucose levels in Type 2 Diabetes (T2D), which is brought about by the insulin resistant state (1). Studies reveal that the *Cebpa* (CCAAT/enhancer binding protein alpha) gene has been recognized as one of the key regulators of gluconeogenic genes. It has distinct regulatory motifs via which it modulates transcription of genes which are essentially involved in the hepatic lipid and glucose metabolism pathways (2, 3). Therefore it is well understood that the *Cebpa* gene is an upstream regulator of hepatic gluconeogenesis and the abundance of C/EBPɑ is an important factor in determining the fate of gluconeogenesis.

Histones are a group of basic proteins that are an integral part of the nucleosome structure. They undergo numerous post translational modifications (PTMs) namely the classical acetylation, methylation, phosphorylation, and ubiquitination and more recently discovered propionylation, malonylation, butyrylation, lactylation and crotonylation (4). There are a number of well established histone marks associated with the expression or silencing of genes. Lysine dimethylation at the 36th position (H3K36Me2) is a marker that is essentially associated with active chromatin and has been observed to upregulate the enhancer activity (5). Evidence has established that histone dimethylation at H3K36 and its consequent demethylation by demethylase Jhdm1a directly regulates hepatic gluconeogenesis (6). This throws light on the fact that *Cebpa* transcriptional activation is authenticated by the presence of histone H3 lysine 36 dimethylation (H3K36Me2) mark on the gene locus. In addition to that, a recent study involving diet-induced obese (DIO) mice showed a very marked increase in the H3K36Me2 methylation in the liver (7, 8). Now, how a histone PTM like H3K36Me2 regulates the transcriptional output of the *Cebpa* gene is just beginning to be understood. Structural and dynamical studies have revealed that behavior of the nucleosome is affected by histone PTMs (9). However, the precise molecular mechanism of H3K36Me2-induced nucleosome dynamics and its role in transcriptional regulation of *Cebpa* is not fully known. Interestingly, while usually acetylation is always associated with activation, methylation of histone can induce both activation and repression depending on where the methylation is present on the histone (10).

Nucleosomes are the structural and functional units of chromatin, consisting of histone octamer and 146 basepairs of DNA in the core (11). They are dynamic chromatin packaging units whose unwrapping and sliding contributes to the availability of the gene for the transcription factors (TFs) to bind and initiate transcription (12). The +1 nucleosome (Nuc+1) positioning near the transcription start site (TSS) can determine gene transcription. It can inhibit (or induce) transcription while positioning very close to (or slightly away from) TSS (≈40–60 basepairs in mammals) (13,14). Recently solved co-structures of Nuc+1 with polymerase or with remodeling enzymes suggest that Nuc+1 can not only unwrap but also reposition during transcription (15–17). Various TFs have been reported to bind throughout the promoter region of the *Cebpa* gene and also to the downstream region of TSS (18, 19). The GTRD or Gene Transcription Regulation Database and JASPAR report TFs that bind to the *Cebpa* gene +1 nucleosome sequences. Thus, these indicate that the Nuc+1 regions may have an eminent role in regulating the expression of genes.

In T2D, the enrichment of H3K36Me2 in the gene region is associated with disease progression (7). While the enrichment of H3K4/H3K9me3 bivalent domains at the *Cebpa* gene locus in adipocytes has been seen to limit the *Cebpa* gene expression by recruitment of factors such as SETDB1 and stalling of RNA-PolII just downstream to the transcription start site (20). These findings infer that histone PTMs have different mechanisms of regulating gene expression which include recruiting other factors at the sites, forming bivalent domains, and repositioning the nucleosome to alter the chromatin accessibility.

One of the ways of capturing and studying the instantaneous structural states of a molecule is through molecular dynamics (MD) simulations. All-atom MD simulations at a larger time scale have been only recently carried out and the structural and dynamic nuances have been reported (21, 22). However, the computational cost is even higher with the incorporated PTMs in the nucleosome systems. Therefore, exploring the effects of PTMs on nucleosome dynamics at a large time scale has called for reduction of the computational cost, which could be achieved using a state-of-the-art forcefield such as SIRAH (South-American Initiative for a Rapid and Accurate Hamiltonian) (23). A recent study of nucleosome dynamics with SIRAH coarse-grained (CG) MD simulations showed “good agreement” with the all-atom nucleosome simulation results, and by now it is a well established fact that post translational modifications of histones affect the nucleosome dynamics and consequently the transcriptional output of genes (9, 24).

In the current work, we study the molecular dynamic mechanism of transcriptional regulation of the *Cebpa* gene by histone PTMs and nucleosome dynamics through long time scale CG-MD simulations. In particular, we uncover how the H3K36Me2 mark might participate in exposure of the TF binding motifs on the nucleosome DNA and could be responsible for increasing the gluconeogenic load in T2D by promoting *Cebpa* transcription. Here, the PTM induces the nucleosome to position away from TSS and causes dynamic unwrapping at the entry DNA that favors transcription. Additionally, we also predict a previously uncharacterised histone mark H2BK108Me2 which may play a role in reversing the structural effects brought about by the H3K36Me2 in altering the transcriptional status of *Cebpa* gene by inducing the nucleosome to encroach the TSS by rewrapping and sliding thereby disfavoring transcription.

## Results

### Effects of histone PTMs on the global conformational dynamics of *Cebpa* nucleosomes

To understand the role of PTMs, particularly involving H3K36Me2 in the transcriptional activation of *Cebpa*, various CG models of nucleosome—five histone PTM combinations and three unique DNA sequences—were modelled and subjected to MD simulations for 5 μs each. Two metabolically relevant TFs namely NR2C2 and GATA4 (25, 26) were found to have binding motifs near to the entry DNA of our modelled nucleosome (Fig 1A). The average structural properties of the mononucleosome systems, after analyzing the trajectories, viz., structural stability (RMSD), compactness (RG), and DNA wrapping (End-End distance) are presented in Fig 1B and Supplementary Fig S2-S5. The average RMSDs of DNA chains lie between 1.5–3 nm and that of the histone globular domains range from 0.8–1.3 nm with standard deviation 0.05–0.3 nm suggesting stable systems without much fluctuation during the entire simulation time (Fig 1B and first and second panel). The average RGs suggest that H3K36Me2+H3K122Ac, H3K36Me2+H2BK108Me2, H3K36Me2+H3K18Me2+H2BK108Me2, and Plus1 (Nuc+1) were found to be structurally more compact with respect to H3K36Me2 (Fig 1B and third panel). To confirm the structural integrity of the histones the individual histone globular domain RMSDs (Supplementary Fig S6), the histone interface distances (Supplementary Fig S7) and the RGs of the histone octamers (Supplementary Fig S8) were assessed for all the three replicates. All the values plateaued around 2μs of the simulations depicting reliable and stable nucleosome structures. To check for the equilibration of the simulations, block convergence analysis was done for RMSD and RG values (Supplementary Fig S9-S11). However, the average End-End distances show that the orientation of the two DNA arms with respect to one another tends to have larger conformational flexibility where they can come as close as ∼2 nm and can be apart as far as ∼16 nm (Fig 1B and fourth panel). The PTMs that have been introduced in the histone chains and an example of the shortest and the largest distances for the Nuc+1 system are shown in Fig 1C and 1D respectively. Thus, the overall nucleosome behaviour containing combinatorial PTMs show structural differences in comparison to no PTMs. The role of each PTM in nucleosome unwrapping and repositioning and how these can affect transcription at a basepair level are explained below.

**Fig 1.**
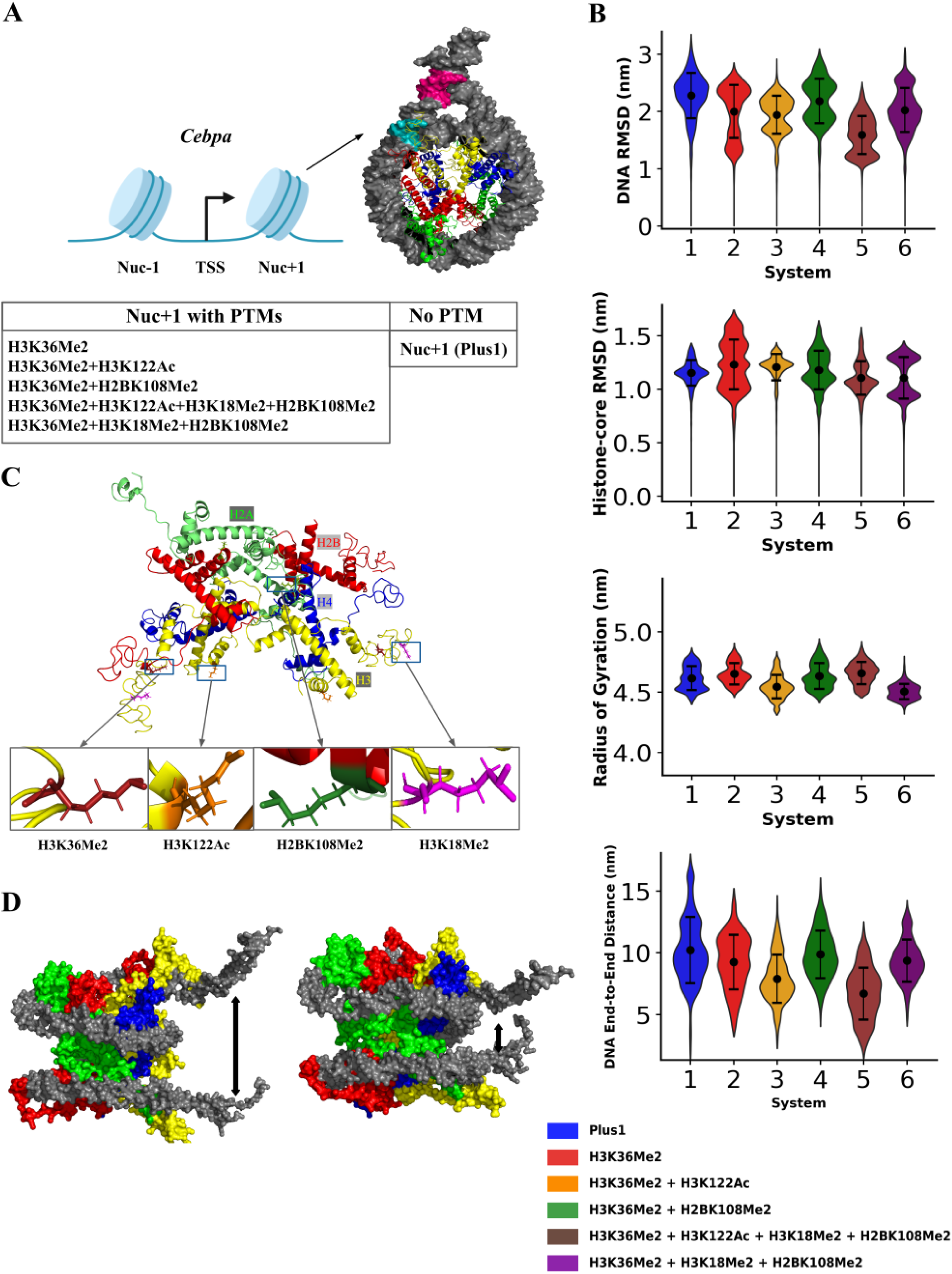
Coarse-grained modeling of *Cebpa* nucleosome with histone PTMs. A) The *Cebpa* locus of interest with TSS, and +1 and -1 nucleosomes. The six mononucleosomes models with and without PTMs are shown in the table below. The binding motifs for TFs NR2C2 (green) and GATA4 (pink) on DNA are shown. B) Average structural properties of the mononucleosome systems obtained from the 5–6*μ*s CG MD simulations viz., structural stability (RMSD) of histone core (first panel) and DNA (second panel), compactness (RG), and DNA wrapping (End-End distance). C) The PTMs are introduced in the histone chains symmetrically. A zoom-view of the PTMs is shown in stick representation. D) Snapshots of CG structures of +1 nucleosome with H3K36Me2 showing the largest (left) and the smallest (right) end-end distances obtained from the simulations. A, C & D) Histones are depicted according to the following colors: H3-Yellow, H4-Blue, H2A-Green, and H2B-Red. DNA is colored as gray.

### H3K36Me2 and co-PTMs of H3K36Me2 promote nucleosome sliding away from TSS

The structural aspect of transcriptional availability can be governed by nucleosome sliding since the loosening or acquisition of DNA at the proximity of the TSS determines whether the transcription factors can bind successfully without any steric hindrance posed by the histones (27, 28). Therefore, we wanted to explore whether there was any sliding in the nucleosome systems, and if any, how they are controlled by the specific PTMs present on the histones. Figure 2A illustrates how the sliding angle was defined and calculated. First we calculated the sliding angle of all the systems (Fig 2B). The sliding angle different from initial angle ≈5–7 degrees indicates that there is either sliding or a combination of sliding and conformational change in histone core octamer. The systems with H3K122Ac tend to show large sliding angles which may suggest deformation in histone core. While the systems Plus1, H3K36Me2, H3K36Me2+H2BK108Me2, and H3K36Me2+H3K18Me2+H2BK108Me2, maintained stable sliding angles throughout the simulation trajectories, suggesting sliding with a stable histone core. However, the direction of the nucleosome sliding was still missing. To address this, the displacement of dyad DNA basepair, towards or away from the TSS was calculated, by defining the entry-exit vector and the direction of DNA wrapping around the histone core. If the dyad basepair shifts towards the entry DNA, the exit DNA is released and vice versa (Supplementary Fig S12A). The collective analysis of all the trajectories for each system revealed that the Plus1 nucleosome maintained a symmetric dyad basepair sliding. Whereas, H3K36Me2 showed the highest dyad basepair shift towards the exit DNA followed by H3K36Me2+H3K18Me2+H2BK108Me2 and H3K36Me2+H3K122Ac+H3K18Me2+H2BK108Me2. This trend was reduced in H3K36Me2+H3K122Ac and H3K36Me2+H2BK108Me2 when compared to the H3K36Me2 system alone (Fig 2C).

**Fig 2.**
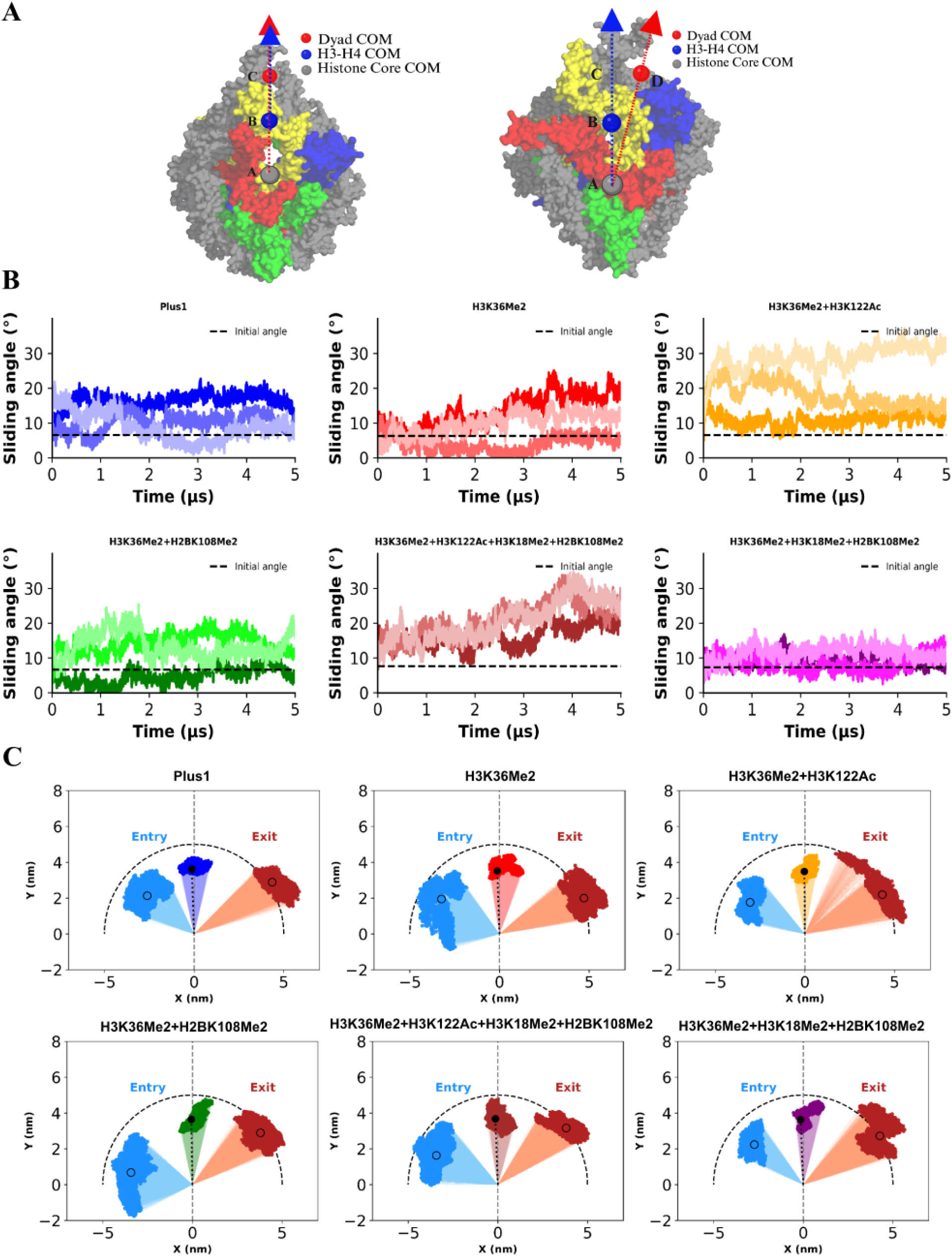
Nucleosome sliding angle and entry-dyad-exit DNA projection. A) The nucleosome sliding angle is defined as the angle between the vector from the centre of mass (COM) of the histone core to the COM of the H3-H4 core and the vector from the COM of the histone core to the COM of the dyad region. The nucleosome color code is the same as in Fig 1. B) Time evolution of the sliding angle of three replicates are shown using three variants of the same color. The dashed line indicates the initial angle. C) Projections of entry, dyad, and exit positions on the X-Y plane where scattered data points are obtained by considering all the three replicates. Shift in dyad position depicts dyad sliding. The initial frame is represented by a black dot with a dashed line. Unfilled circles in the entry and exit regions are the mean values.

### Nucleosome containing H2BK108Me2 promotes TSS-proximal wrapping

The TSS-proximal DNA unwrapping is one of the primary determinants of TF binding and transcription initiation. The DNA end-end distance (Fig 1B) and entry-exit projection (Fig 2C) indicate that nucleosome unwrapping dynamics can be affected by PTMs. However, the extent of unwrapping can not be definitively known. Therefore, we calculated the basepair DNA unwrapping on the entry and exit side of all our nucleosome systems (Fig 3). H3K36me2 containing nucleosomes exhibited a greater entry DNA unwrapping indicating a TF permissive state. Whereas, the H2BK108Me2 containing systems maintained low basepair unwrapping values, comparable to the Plus1 nucleosome. This suggested that H2BK108Me2 may favor a more stable and wrapped conformation near the TSS, restricting the spontaneous DNA opening which can influence its accessibility by the TFs.

**Fig 3.**
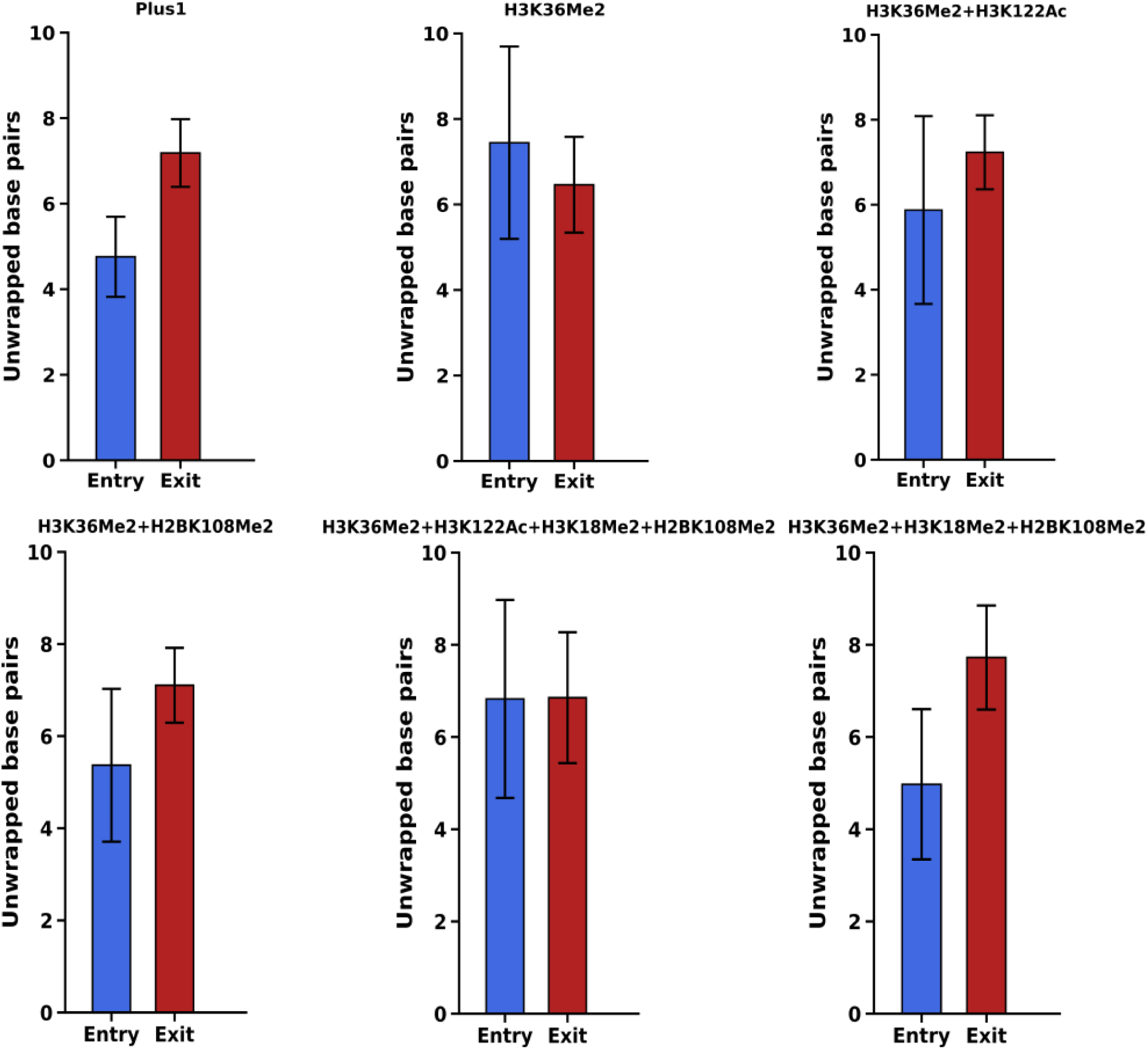
Unwrapping lengths of entry and exit DNA. Bar plots show the average number of unwrapped basepairs on the entry and exit side of the nucleosomes containing combinatorial PTMs on *Cebpa* +1 nucleosome. All three replicates for each system have been considered and the error bars indicate the variability among them.

### *Cebpa* Nuc+1 can exist mainly in two distinct conformational states of nucleosomal DNA

The DNA conformational variability across the nucleosome systems were quantitatively compared by performing principal component analysis (PCA) of the DNA throughout the simulation trajectories (Supplementary Fig S13A), followed by K-means clustering using silhouette scores (Fig 4A). The resulting PCA scatter plot revealed two distinct structural states of DNA (Fig 4B and Supplementary Fig S13B). Cluster 0 consisted of partially unwrapped DNA with the entry side more unwrapped than the exit side (Fig 4C). About 46% of the H3K36Me2 DNA conformations and 31% of the H3K36Me2+H3K122Ac conformations belonged to this cluster, whereas only 25% of the spatial structures of H3K36Me2+H2BK108Me2 was in cluster 0. Cluster 1 showed a more tight DNA wrapping around the histones displaying a more canonical nucleosome structure (Fig 4D). The H2BK108Me2 histone mark containing systems majorly occupied cluster 1, with 75% of the structures of H3K36Me2+H2BK108Me2 and a full 100% of H3K36M2+H3K18Me2+H2BK108Me2, indicating that H2BK108Me2 might be important in the events of nucleosome wrapping. Both the clusters are associated with the DNA dyad sliding away from the transcriptional start site of the nucleosome (Fig 4E).

**Fig 4.**
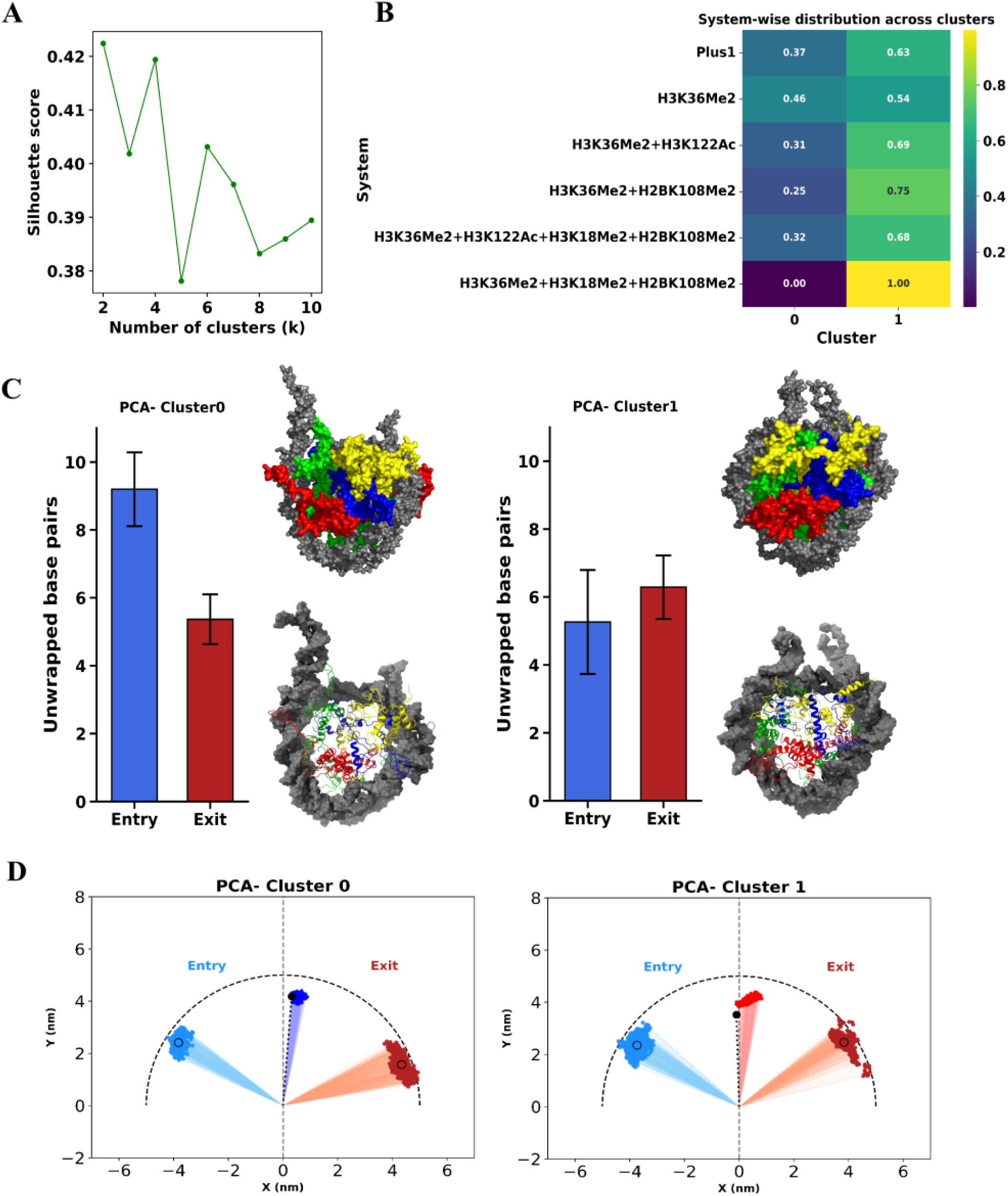
PCA of nucleosomal DNA trajectory. A) Silhouette score analysis to determine optimal number of clusters, here k=2, suggesting two distinct conformational states of the nucleosomal DNA. B) Cluster composition heatmap exhibiting the fractional contribution of each system across the two clusters. C) Unwrapping lengths of entry and exit DNA of Cluster 0 (left panel) and 1 (right panel) nucleosome structures. Top panels are coarse-grained representation, and bottom panels are all atom representation. The nucleosome color code is the same as in Fig 1. D) Entry-dyad-exit DNA projection on the X-Y plane using 500 samples obtained from the PCA clusters. The initial frame is represented by a black dot with a dashed line. Unfilled circles in the entry and exit regions are the mean values.

### TSS-proximal DNA and histone contacts of *Cebpa* Nuc+1 are regulated by histone PTMs

Histones can contact DNA either by histone globular domain or histone tails or combination of both. To assess the contribution of histones in the nucleosome stability, contact distance map was computed. We simulated five different combinations of histone PTMs associated with T2D on the +1 nucleosome of the *Cebpa* gene. An initial observation was that PTMs induced changes in DNA-histone interaction patterns across all cases which are revealed by the DNA-histone contact maps and heatmaps (Supplementary Fig S14A-F & Fig S15A-F). Moreover, trajectory visualization clearly demonstrated that histone tails play an essential role in mediating DNA-histone contacts within the nucleosome. To substantiate this finding, DNA-histone globular domain contact maps and heatmaps were subsequently constructed (Supplementary Fig S16A-F & Fig S17A-F respectively). We were further interested in finding out how each of the histone tails (H3, H4, H2A, and H2B) and the histone globular domain participate in the interactions with DNA (*Cebpa* gene) and how this pattern changes due to the varying PTM marks that are carried by the histones. In order to study that, the histone components were divided accordingly and the mean contact of each component against the DNA basepairs were calculated. Each PTM system exhibited distinct contact footprints (Supplementary Fig S18A-C), with the core maintaining the highest contact frequencies but differing among systems. H3K36Me2+H2BK108Me2 and H3K36Me2+H3K18Me2+H2BK108Me2 showed the most consistent contacts frequencies of the entry DNA or the site close to the TSS, of the nucleosome (around basepairs -93 to -70), to the histone globule domains, followed by Plus1, H3K36Me2 and H3K36Me2+H3K122Ac containing nucleosome systems. The replicates also showed similar histone footprints. This observation further helped us conclude that H3K36Me2+H2BK108Me2 modification closes the nucleosome on the TSS side by increasing DNA-histone interaction at the entry point. Therefore, it’s worth mentioning here that the spatial heterogeneity in the loss of contacts that can be observed throughout the trajectories especially at the nucleosome edges is suggestive of PTM-driven local and global weakening of DNA-histone interactions.

### *Cebpa* Nuc+1 can exist mainly in three distinct conformational states based on histone-DNA interactions

To visualize the influence of the various histone modifications (single and combinatorial), Uniform Manifold Approximation and Projection (UMAP) was applied to high dimensional DNA-histone distance data. The UMAP projection of DNA-histone distances (Supplementary Fig S19A) revealed formation of very clear and spatially well separated clusters according to histone modification states in the 2D embedding space. This is indicative of the fact that different post translational signatures pose unique chromatin conformations. To further analyse which of the clusters occupied isolated chromatin states, K-means clustering were performed following the silhouette score calculations (Fig 5A). Consequently, three distinct clusters formed, under which we could place the PTM systems (Fig 5B and Supplementary Fig S19B). The most striking observation was that the system with H2BK108Me2 restored PTMs—H3K36Me2+H2BK108Me2 and H3K36Me2+H3K18Me2+H2BK108Me2—were populated in cluster 0 with 84% and a full 100% of the conformations respectively. This observed cluster preference indicated that H2BK108Me2 promotes a distinct DNA-histone interaction pattern and locks the chromatin in an unique conformational state. Further investigation was done on the clusters identified by UMAP to understand whether they correspond to unique DNA accessibility states of the nucleosomes. Each cluster exhibited distinct DNA conformations portrayed by different extent of the entry side DNA breathing. Cluster 0 depicted the greatest entry DNA unwrapping of ≈9–10 basepairs (Fig 5C and left panel). Whereas, Cluster 1 displayed a more canonical nucleosome structure with lesser asymmetry between entry and exit DNA unwrapping (Fig 5C and right panel). Cluster 2 represented the most closed state of the entry DNA among all (Fig 5C and bottom panel). Cluster 1 and 2 are visited by Plus1 system with 47% of the structures in Cluster 2, H3K36Me2 with 33%structures in Cluster 2 depicting mild decompaction to a more open nucleosome, nearly comparable to H3K36Me2+H3K122Ac with 47% of the configurations in Cluster 2.

**Fig 5.**
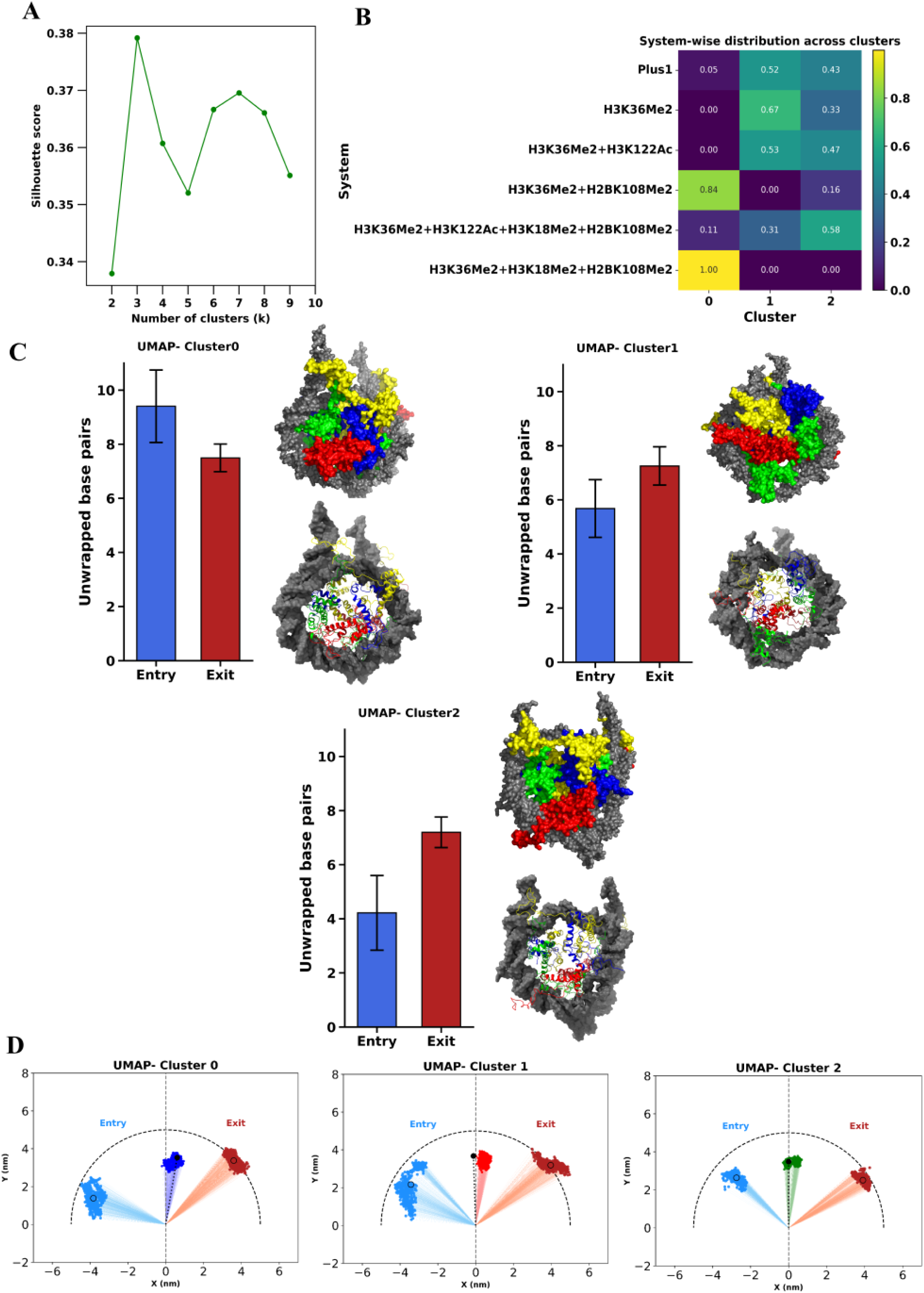
UMAP of histone-DNA interaction trajectory. A) Silhouette score analysis to determine the optimal number of clusters ranging from 2-10, showing the highest value at k=3 i.e., the optimal clustering solution. B) The fractional contribution among the 3 clusters of each PTM containing nucleosomes in the cluster composition heatmap. C) Unwrapping lengths of entry and exit DNA of Cluster 0 (left panel), 1 (right panel), and 2 (bottom panel) nucleosome structures. Top panels are coarse-grained representation, and bottom panels are all atom representation. The nucleosome color code is the same as in Fig 1. D) Entry-dyad-exit projection on the X-Y plane using 500 samples obtained from the UMAP clusters. The initial frame is represented by a black dot with a dashed line. Unfilled circles in the entry and exit regions are the mean values.

### A mechanism of H2BK108 dimethylation-dependent regulation of *Cebpa* +1 nucleosome accessibility in gluconeogenesis

Under normal conditions, the *Cebpa* +1 nucleosome maintains a stable entry DNA wrapping occluding the TSS-proximal region supporting a basal transcription maintenance of the gene (Fig 6, left panel). In the altered metabolic circumstances such as Type 2 Diabetes, the H3K36Me2 histone PTM mark is elevated. Our model shows a higher degree of nucleosome sliding away from the TSS complemented with entry DNA unwrapping in the H3K36Me2 system (Fig 6, middle panel). This suggests increased accessibility to the transcription factor binding motifs near the TSS and promotion of *Cebpa* transcription. Restoration of the H2BK108Me2 histone mark attenuates entry DNA unwrapping and decreases the level of nucleosome sliding as compared to H3K36Me2 mark alone, which decreases the TF accessibility to the TSS-proximal DNA and suppresses *Cebpa* transcription (Fig 6, right panel).

**Fig 6.**
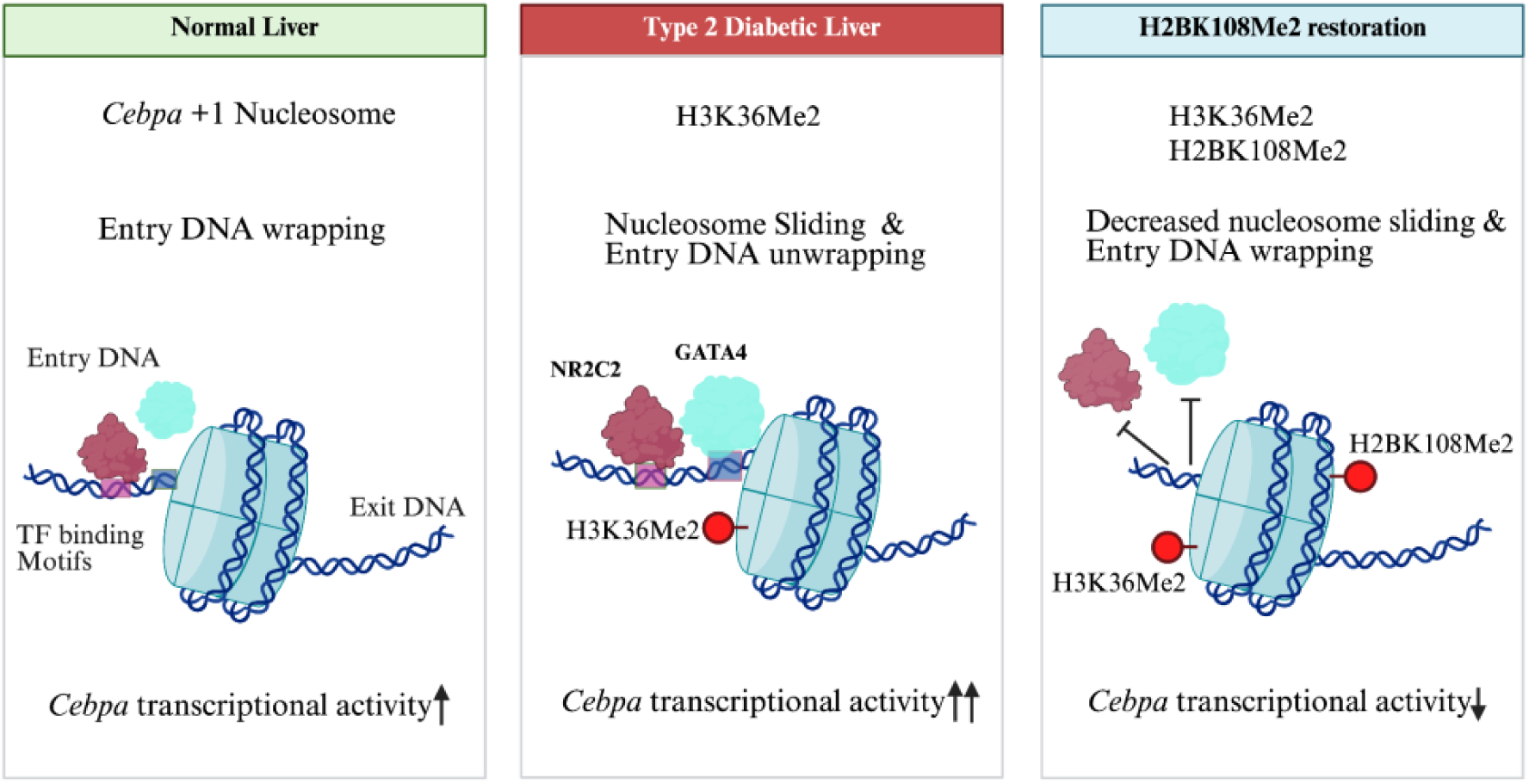
A model of H2BK108Me2-mediated transcriptional regulation in the *Cebpa* +1 nucleosome. The schematic illustrates the effects of histone PTMs on nucleosome dynamics of the *Cebpa* +1 nucleosome near its TSS. (Left panel) The DNA wrapping near the TSS under normal conditions. (Middle panel) In T2D, increase in H3K36me2 mark causes TSS proximal DNA unwrapping making the gene more gullible to transcription. (Right panel) H2BK108me2 when restored, reduces nucleosome sliding and promotes entry DNA wrapping to decrease the entry DNA accessibility and suppress *Cebpa* transcription.

## Discussion

Nucleosomes are highly dynamic structures undergoing unwrapping, rewrapping and repositioning or sliding which are the key aspects of gene expression regulation (29). Here, the effect of histone PTMs on the nucleosome dynamicity in the context of T2D has been presented. This study has explored how the histone H3K36 dimethylation mark could be associated with increasing the transcriptional availability of *Cebpa* gene and might play an eminent role in increasing the downstream gluconeogenic flux purely by altering the structural integrity of the *Cebpa* +1 nucleosome, adding to the present scenario that dimethylation causes gluconeogenic surge.

The MD simulations of nucleosomes have emerged to be a powerful tool to study the real-time changes in the interaction patterns within the structure (22, 30). All-atom MD simulations appear to be an effective complement to the experimental studies going on in the field. However the computational cost of performing them at large time scales is something that we are trying to overcome collectively. This problem has been resolved to a certain extent by using coarse-grained (CG) models for simulating the large systems using forcefields such as MARTINI (31) and SIRAH (23). These have enabled us to run MD simulations for large systems at microsecond time scales, but they come with certain compromises as well. The limitations of SIRAH coarse-graining currently include loss of atomic-level precision, overrepresentation of peptide folding and exclusion of parameterization of biological molecules due to their vast variations (32). So, its usage is highly dependent on what biological question we are trying to answer (23). Although nucleosomal DNA unwrapping and sliding are events which occur at higher temporal resolutions, our coarse-grained MD simulations can capture the dynamic partial unwrapping and rewrapping like events and dyad sliding and DNA shift around the histone core. This can be attributed to the coarse-graining of our models and as compared to the all-atom models, the CG models are two to three times faster. This happens effectively due to the combined influence of fewer degrees of freedom, faster sampling because of the smoothened energy landscape and higher integration time steps (33). Thus, the 5 microsecond MD simulations reported here have revealed the clear distinction in DNA-histone interaction footprints with the changing PTM marks associated with the nucleosome histones.

The DNA-histone interactions and their deviations are prompted by the PTMs which are differentially expressed in the diseased condition. Single molecule tweezer experiments and both CG and atomistic simulations have shown that even confined nucleosome sliding was sufficient to expose or occlude nearby TF (such as Egr-1) binding sites, and a shift of only ≈1–2 basepairs can alter the rotational exposure for the pioneer factor motifs such as Sox2 and Oct4 which consequently change the TF affinity, its mode of binding and cooperativity (34, 35). Our simulation data not only replicates the trend of previous findings about H3K36Me2 increasing the transcription of the *Cebpa* gene but also throws light upon the mechanism by which it may operate (6).

The events associated with the changing dynamics of a nucleosome system are dependent on multiple factors such as histone PTMs, DNA methylation, and the DNA sequence itself (36). Nucleosome repositioning can occur via the DNA sliding along the histones and can be achieved either through loop propagation or twist diffusion (37). While the loop propagation depends on ATP-dependent remodelers, the twist diffusion-mediated nucleosome sliding is primarily driven by DNA sequence-dependent properties and thermal fluctuations. The mechanism of DNA sequence-dependent nucleosome sliding is closely related to the deformation variables of DNA such as twist, roll, tilt, shift, and rise (37). Moreover, the sliding has been reported to be directly dependent on the binding energy of the histone core to a given DNA sequence (38). In line with these, our simulation data suggest that the observed nucleosome sliding is due to the twist diffusion mechanism.

Although sliding is an important phenomenon for creating nucleosome depleted regions, DNA unwrapping has been found to be one of the primary determinants of TF binding on its respective motifs (39, 40). The integration of our results from the trajectory analysis and the structural visualization confirmed the sliding of H3K36Me2-nucleosomes in the direction of the TSS-proximal entry DNA releasing more DNA near the TSS. Further, the increased TSS-proximal DNA unwrapping increases its possibility to be available for the plausible transcription factors such as NR2C2, GATA4, HNF family or pioneer factors such as CEBP family to bind (18, 25, 26, 41). This study however particularly focuses upon the effects of the associated histone PTMs and not the DNA sequence on *Cebpa* +1 nucleosome dynamics. The possibility that these results could be highly specific for this gene can not be ruled out.

H2BK108Me2 is one of the histone PTM marks found to be downregulated in the DIO mice liver (7). Our study suggests that the structural effects posed by H3K36Me2 can be rescued via nucleosome sliding in the opposite direction towards the TSS with the restoration of the PTM mark H2BK108Me2 on the +1 *Cebpa* nucleosome. These observations may be largely due to PTMs since the only variable throughout our models and simulations was the combination of different PTMs on the nucleosomes. This study provides mechanistic insights into chromatin regulation paradigms and reveals that various PTMs and their combinations are indeed capable of exerting differential modulations in nucleosome structure and chromatin accessibility at the TSS. This in turn can regulate the downstream transcriptional output and metabolic status of a cellular system.

There are no previous reports as of yet about how H2BK108Me2 participates in the regulation of chromatin accessibility. Our computational experiments and results have directed us to infer that H2BK108Me2 may be associated with nucleosomal closure towards the TSS in *Cebpa* +1 nucleosome and thus it can be one of the important target methylation sites for the treatment and management of T2D.

### Conclusion and Future prospects

This study comprehensively demonstrates the probable mechanism of gene expression regulation due to nucleosome dynamicity and reveals that histone PTMs tend to regulate transcriptional flux of *Cebpa* gene, and also shows that their effects can be regulated by other counteracting histone PTMs. This study predicts that the presence of H2BK108Me2 at the *Cebpa* +1 nucleosome may repress its expression by repositioning the nucleosome at TF binding sites. Thus, H2BK108Me2 has emerged as one of the possible target histone PTMs which holds great potential towards the reduction of gluconeogenic load in T2D. Our hypothesis and results shape the basis of future experiments to study H2BK108Me2 as a useful target for development of drugs and management of the disease. Our predictions can be experimentally validated by perturbing H2BK108Me2 histone mark followed by examining the changes in chromatin accessibility and nucleosome occupancy using MNase-seq, ChIP seq or ATAC-seq based assays. Finally, our study provides the chromatin basis of T2D and an explanation of the impacts of structural chromatin regulation on metabolic cascade of a system such as the gluconeogenesis pathway.

## Materials and Methods

### Reagents and Tools

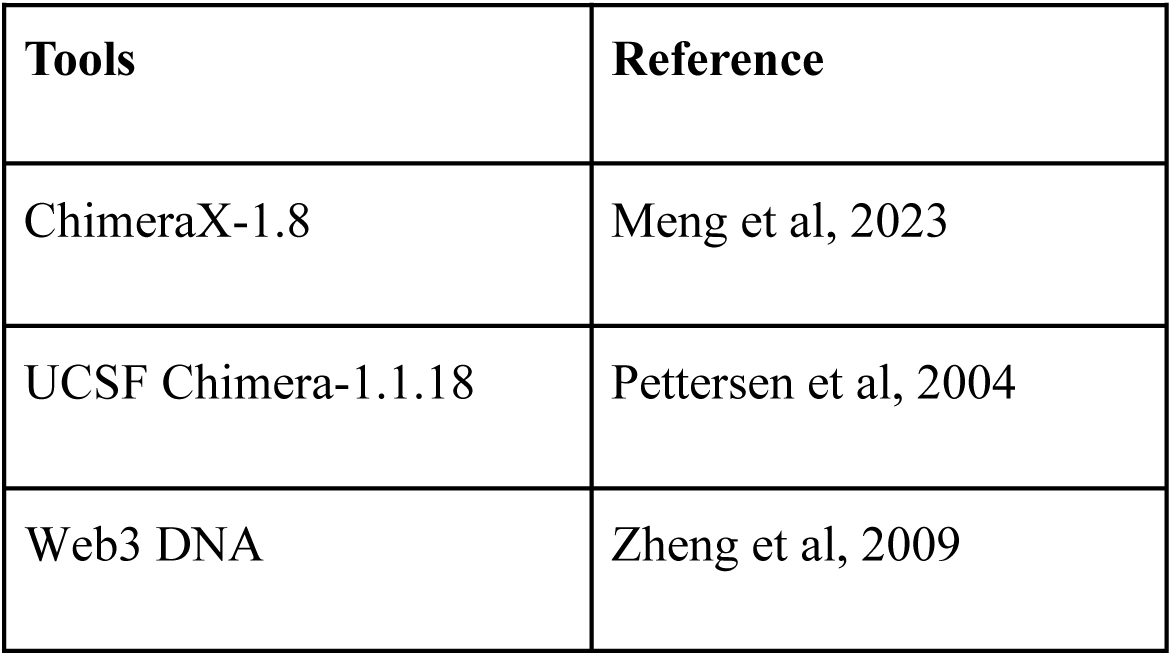
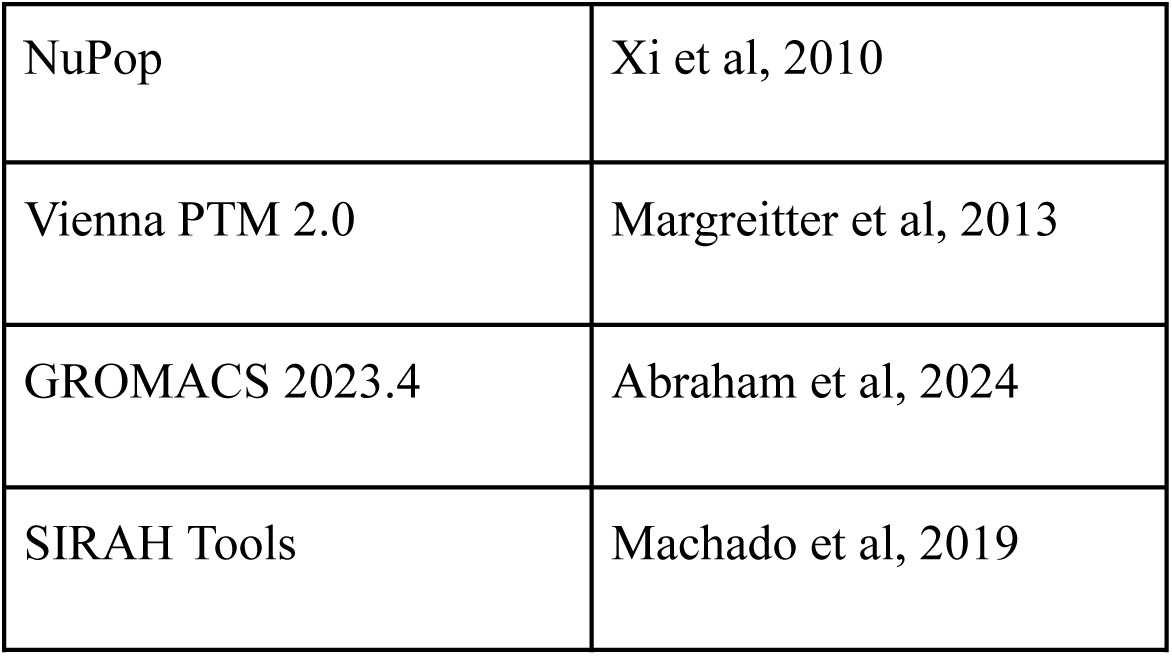

Reagents and Tools:
| Tools | Reference |
| --- | --- |
| ChimeraX-1.8 | Meng et al, 2023 |
| UCSF Chimera-1.1.18 | Pettersen et al, 2004 |
| Web3 DNA | Zheng et al, 2009 |
| NuPop | Xi et al, 2010 |
| Vienna PTM 2.0 | Margreitter et al, 2013 |
| GROMACS 2023.4 | Abraham et al, 2024 |
| SIRAH Tools | Machado et al, 2019 |

### Selection of gene and histone post-translational modifications

For the study, *Cebpa* gene was selected due to its already established role at hepatic glucose metabolism as a critical transcriptional regulator influencing gluconeogenic pathways and metabolic homeostasis in the liver. Evidence from Pan *et al.,* 2012 demonstrates that H3K36Me2 is abundantly present at the *Cebpa* locus and is implicated in transcriptional control relevant to gluconeogenesis (6). Then the PTMs associated with T2D were focused upon. From the findings by Nie *et al.*, 2017 the H3K36Me2 emerged as a notably upregulated histone mark in DIO mice liver (7). This suggested its probable involvement in the regulation of metabolic genes under diabetic conditions. Although these PTM profiles were derived from mice, histones and the modification sites are highly conserved between mice and humans, enabling direct mapping of these PTMs onto human nucleosomes, and effectively disbursing their individual or combinatorial effects (42, 43). Thus, our analysis focuses on the structural and biophysical impact of these modifications while acknowledging potential species-specific chromatin differences. Therefore, these two strands of evidence were integrated and sewn. *Cebpa*, a T2D related relevant hepatic gene, and H3K36Me2, an upregulated epigenetic mark in altered metabolic condition were taken into account and the computational experiments were performed to uncover the relationship between histone PTMs and nucleosome dynamics modulations in setting of T2D.

### Construction of nucleosome models with native human DNA sequence of *Cebpa* gene

Eight nucleosomal systems were modelled with 186 basepairs of DNA, 146 basepairs for nucleosome core and 20✕2 linker basepairs on either side. The systems were constructed with the DNA from the Plus 1 region of the *Cebpa* gene (i.e., they are Nuc+1 nucleosomes). All the nucleosomes were constructed over the X-ray diffraction structure PDB ID 3AFA (44) and the DNA sequence for probable nucleosome positions were determined using the NuPop program (45). Additionally we analysed the MNase-seq dataset GSM3718063 (46) to determine the +1 and –1 nucleosome positions on the *Cebpa* gene (Supplementary Fig S1). The transcription start site was determined using the Genome Data viewer webserver on GRCh38/hg38 (accessed on: 25/01/2025). The sequences for the *Cebpa* gene were retrieved from UCSC genome browser (hg38) (accessed on: 25/01/2025). The missing histone residues were modelled using modeller in ChimeraX-1.8 (47) and the *Cebpa* +1 DNA were constructed using Web3 DNA program (48) (accessed on: 25/01/2025) and then incorporated into the existing structure by replacing the DNA in the 3AFA.pdb using UCSF chimera (version:1.1.18) program (49). All the histone tails were left nearly at their original positions they had assumed, only the portions of the tails which were representing clashes/contacts were adjusted spatially to avoid any kinds of clashes. The desired PTMs were incorporated using the Vienna PTM 2.0 program (50) symmetrically on both copies of each kind of histone chain (accessed on: 26/01/2025).

### Molecular Dynamic Simulations Setup and Pipeline

All the nucleosomes were modelled using the atomic coordinates of the nucleosome with PDB ID 3AFA in an all-atom ensemble using UCSF Chimera (version 1.1.18) and web3 DNA program. Then the structures were protonated using UCSF chimera. The coarse-grained mapping of the nucleosome systems with desired PTMs were done using SIRAH tools with the parameters provided by the SIRAH v2.1 force field (23). SIRAH offers an array of PTM parameters included in their mapping directory of sirah.ff, which were the source for the corresponding residue topologies of PTMs (lysine dimethylation and lysine acetylation) ensuring the compatibility with the original SIRAH 2.1 parametrization scheme. Once the structures were coarse grained they were taken ahead through different steps of MD simulations using GROMACS. All the simulations were carried out in replicates of three for each of the five systems with GROMACS version 2023.4 on GPU accelerated nodes (51).

### MD simulations Protocol

1. The nucleosomes were placed at the centre of a cubical box and vacuum minimization was carried out with the steepest descent method with a maximum of 50,000 steps until the maximum force reached 500 kJ/mol/nm.
2. After the vacuum minimization, SIRAH solvent molecules WT4 were added to the system followed by the addition of monovalent NaCl at a concentration of 0.15 M. Another round of minimization was carried out after adding solvent and ions with the same specifications mentioned earlier.
3. NVT equilibrations were performed using the leap-frog integrator for a maximum of 500,000 steps (1ns). Temperature annealing was performed over the course of the equilibration from 40 K to 310 K using a single-stage ramping across ten time points. Nonbonded interactions were handled using the Verlet cutoff scheme (52). Particle Mesh Ewald (PME) method (53) with a Fourier grid spacing of 0.12 nm was employed for the long range electrostatic interactions. Periodic boundary conditions in all dimensions were used. The system was coupled with a modified Berendsen thermostat (V-rescale) (54) with a relaxation time of 0.1 ps.
4. The NPT equilibration had similar parameters to that of the NVT. Parrinello–Rahman barostat (55) was used to maintain the pressure at 1 bar with a coupling constant of 2.0 ps and isotropic pressure scaling. The compressibility was set to 4.5×10⁻⁵ bar⁻¹, approximating that of water.
5. Production run was carried out on the NPT ensemble, with pressure at 1 bar for a time scale of 5 microseconds for each model system with the integrator time step of 10fs and the output was saved every 1ns. Bond constraints were maintained using LINCS algorithm (56).

### Analysis of the MD trajectories

In order to establish the role of the particular PTMs in controlling the nucleosomal behaviour and how the modulation of the structures in turn determine the transcriptional output, we took to explore the changes in the histone-DNA interaction patterns, the nucleosome sliding angle and the nucleosome sliding direction due to the PTMs on histones. Since, our sole focus was on the PTMs, no other factors were considered for these analyses. All three trajectories for each system were taken into account for the analysis. For the quantification of the parameters mentioned, python codes were developed (v3.12) (57) and the MDAnalysis package was used (58, 59). The interaction distance between histone and DNA were taken as 1nm in each case. Apart from that, functions like gmx_rms, gmx_gyrate, gmx_distance functions form the GROMACS software were used to analyse the Root Mean Square Deviation (RMSD), Radius of Gyration (RG), and end-to-end (End-End) distance between the entry and exit DNA ends in order to understand the overall and local stability and dynamics of each model system. The end to end distance of the DNA was calculated between the phosphate atoms of the first and the last basepairs of 146 basepairs of DNA (the linker DNA was excluded) with the gmx_distance function. The sliding angle was calculated as the angle between the two principal axes that we have defined below:

1. The vector from the centre of mass (COM) of the histone globular domains to the COM of the dyad basepair.
2. The vector from the centre of mass (COM) of the histone globular domains to the COM of the H3-H4 globular domains.

The change in this angle was captured with the progress of the simulation and translated into the essence of nucleosome sliding. The shift of the dyad basepair by any given angle implied the displacement of DNA along the histone. A similar study by Niina *et al.,* 2017 was carried out to study nucleosome sliding (27). Our sliding angle and direction calculations were validated using their method. The entry and exit DNA basepairs were defined by excluding the linker DNAs. Unwrapping was calculated by counting the number of DNA entry and exit DNA basepairs that lost contact with the histone globular domain, following the contact criteria in the work of Armeev *et al.,* (22).

## Acknowledgement

We acknowledge the financial support provided by IIT Mandi Seed Grant (Grant Nos. IITM/SG/SS-DDP-KH-RKO/123) and high-performance computing resources provided by Param Himalaya at IIT Mandi implemented by CDAC and funded by the Government of India.

## Author Contribution

Ananya Chandra: Conceptualization, Experiment designing and carrying out the experiments, Data Curation, Analysis, Writing-original draft, Writing-review and editing. Kharerin Hungyo: Conceptualization, Experiment designing, Writing-review and editing, supervision. All authors contributed to reviewing and editing the manuscript. All authors reviewed, checked, and approved the final manuscript.

## Data Availability

**Structure files and trajectories** are available in Zenodo (https://doi.org/10.5281/zenodo.17736189).

## Supplemental information

**Table S1.**
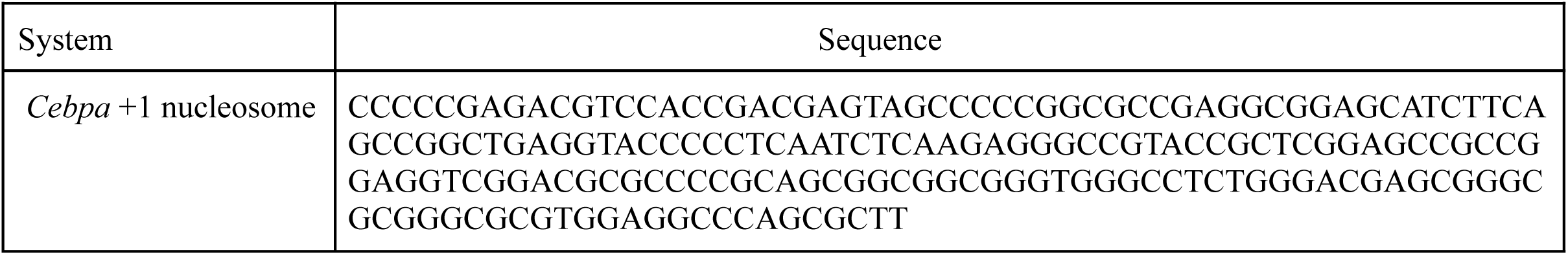
The genomic DNA sequence of *Cebpa* +1 nucleosome.

**Fig S1.**
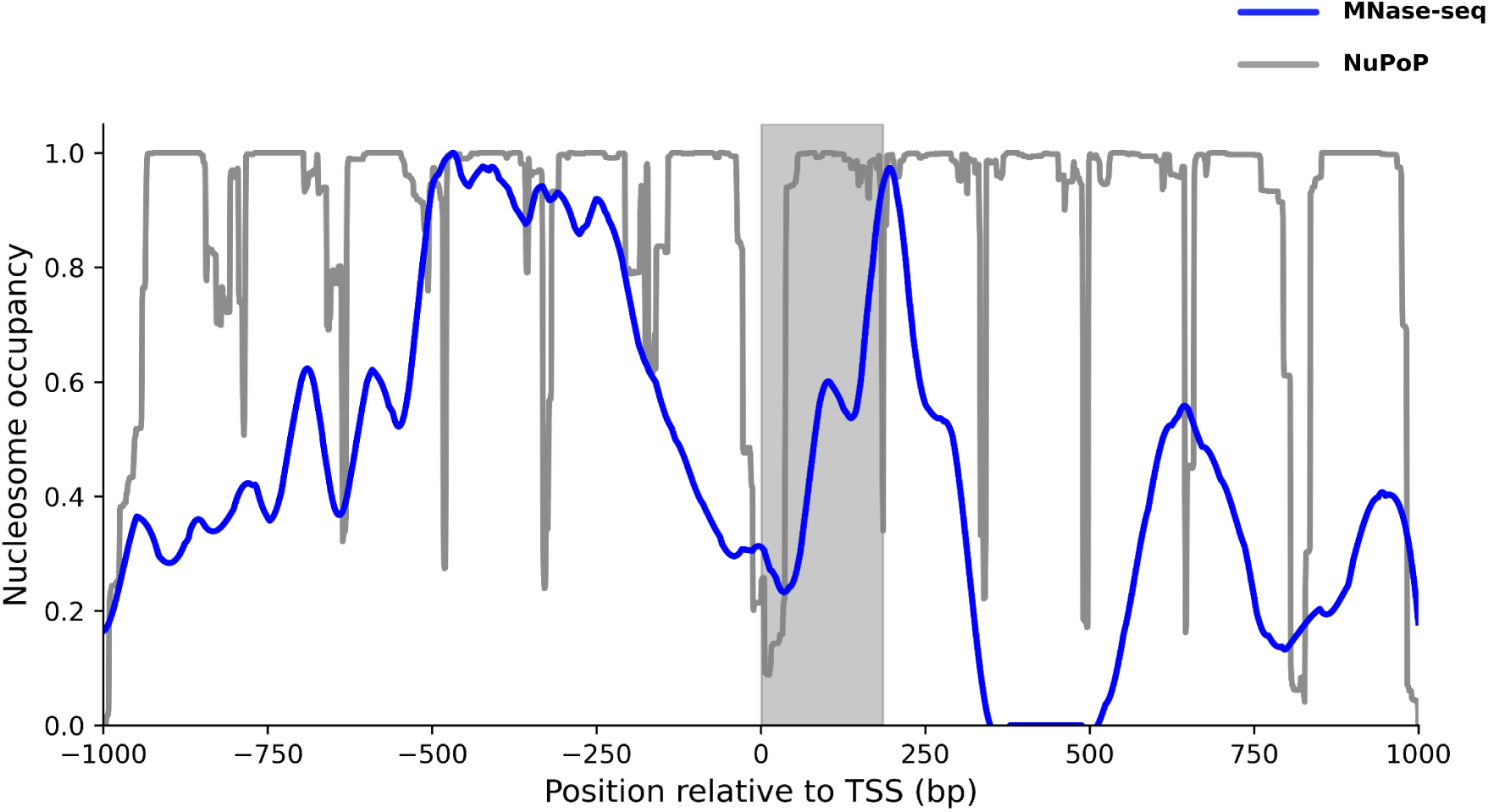
*Cebpa* +1 nucleosome located on the genome from MNase-seq dataset GSM3718063.

**Fig S2.**
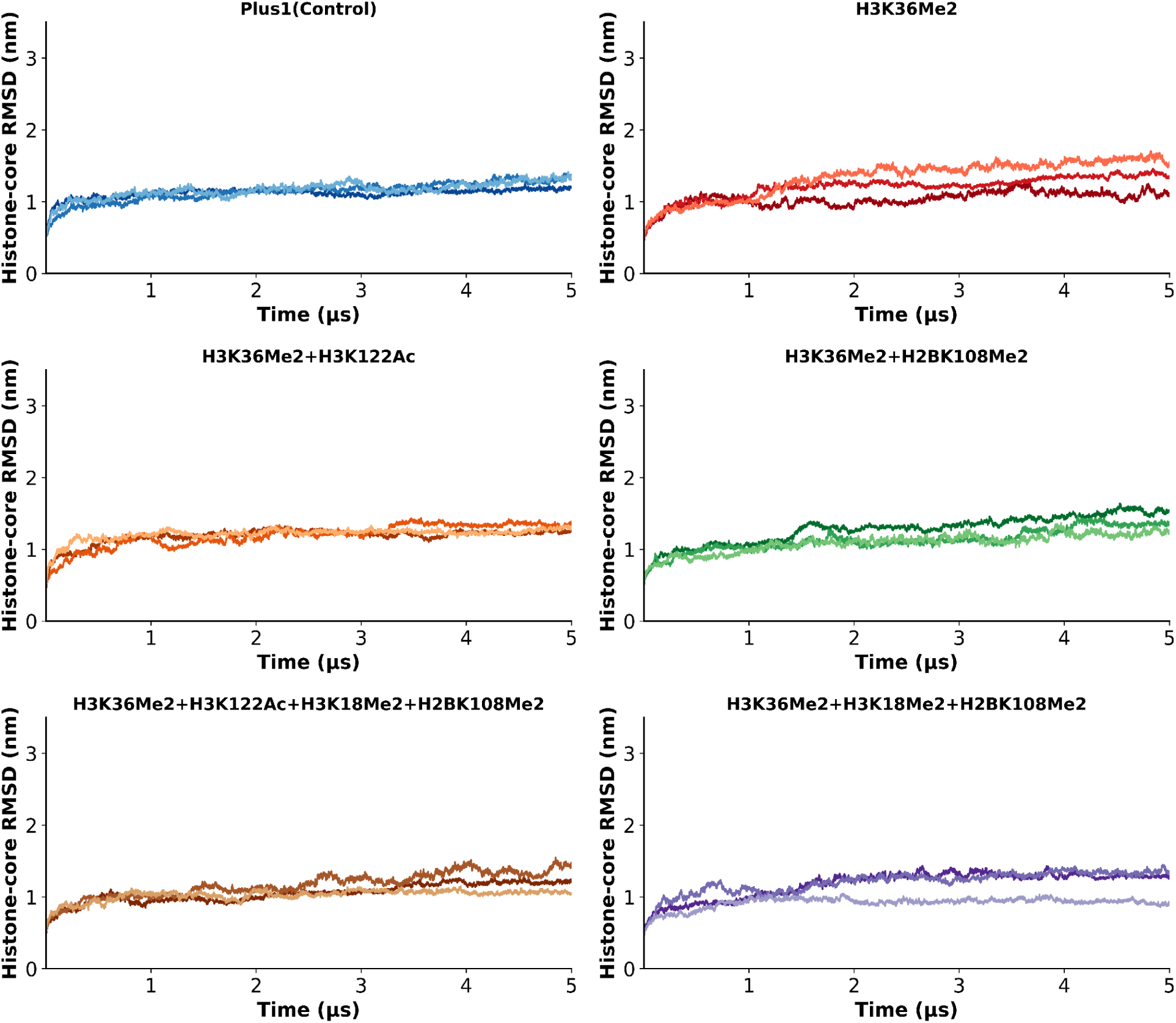
Depicts time evolution of RMSD of histone globular domain. The three replicates are shown using three variants of the same color.

**Fig S3.**
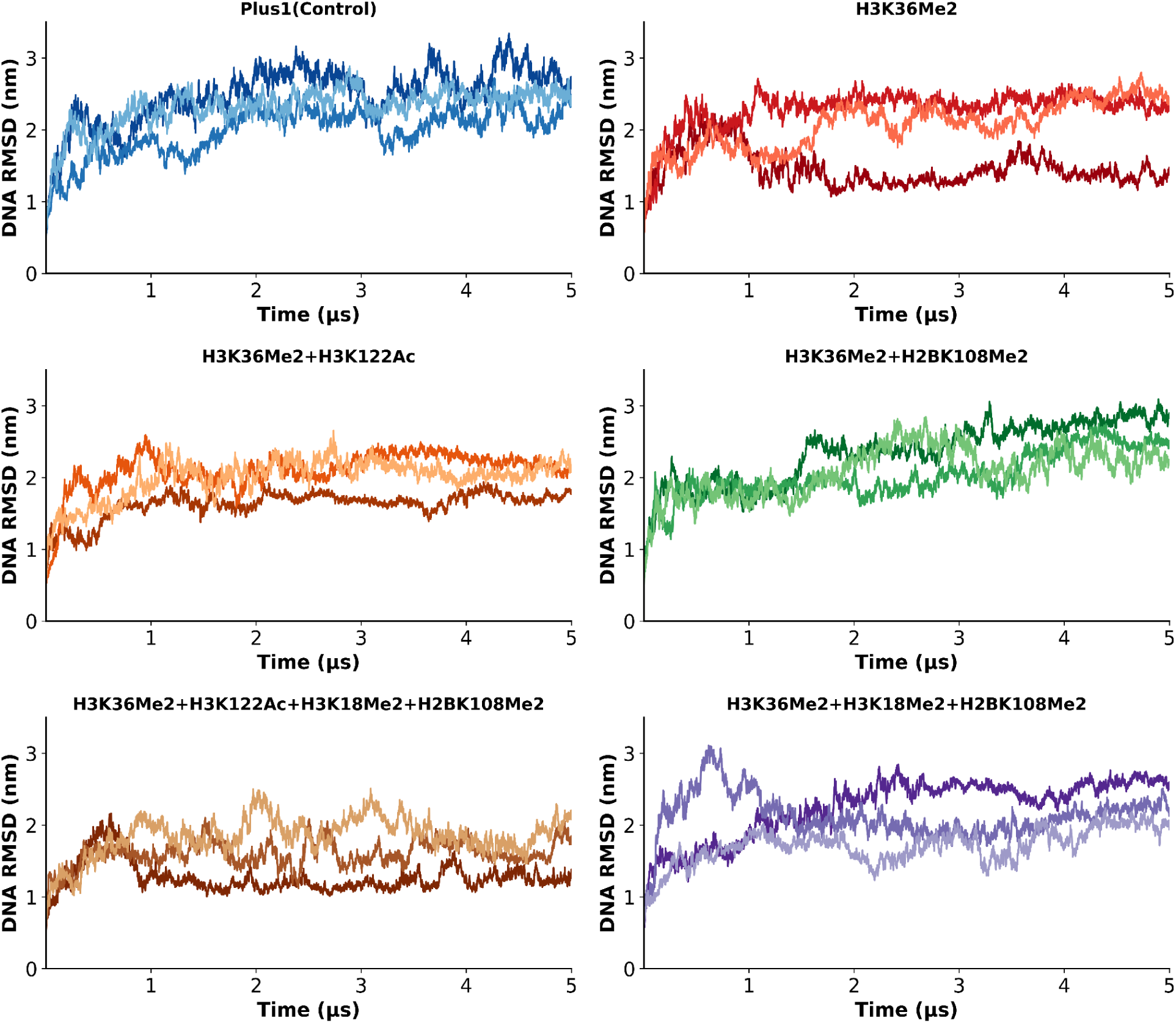
Depicts time evolution of RMSD of nucleosome DNA. The three replicates are shown using three variants of the same color.

**Fig S4.**
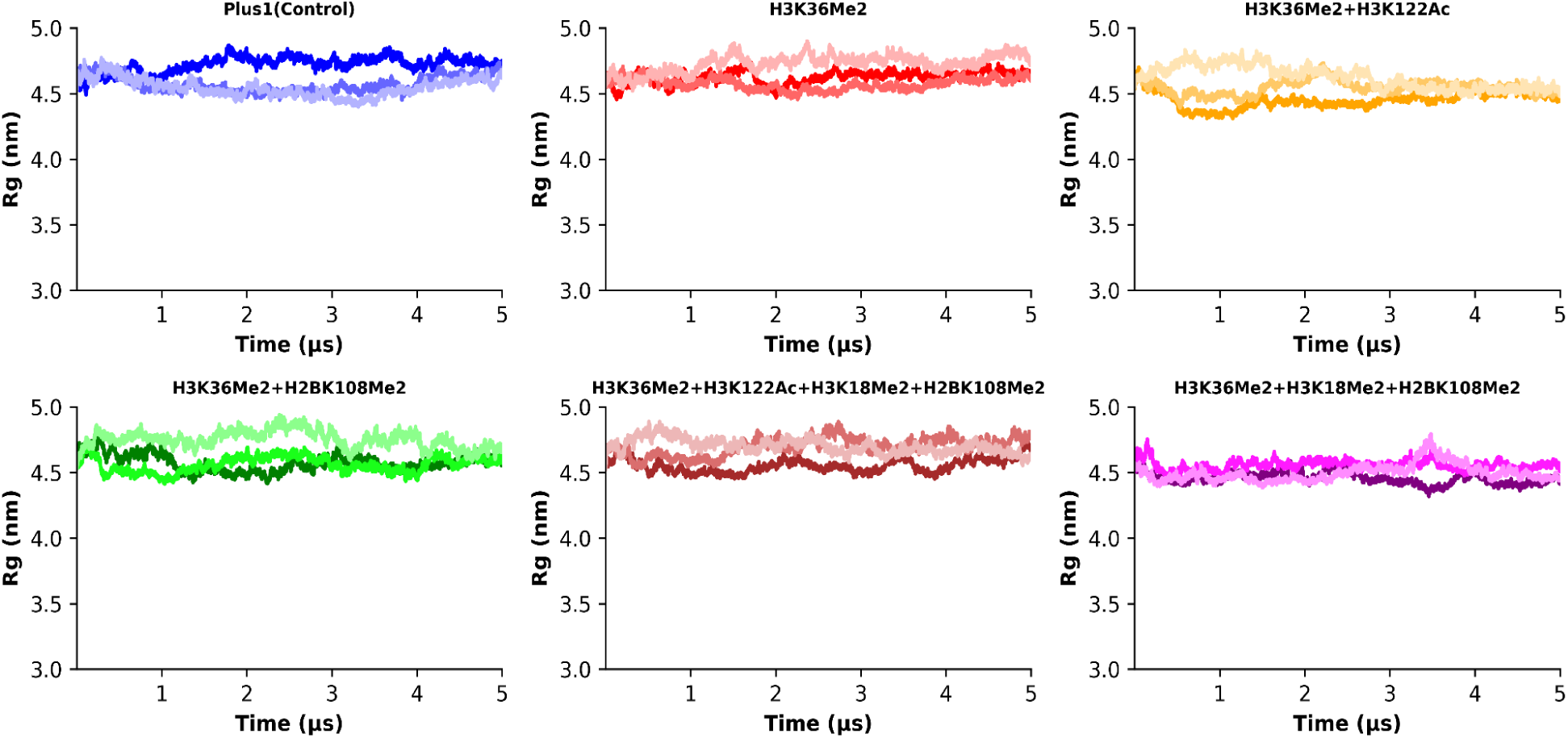
Depicts time evolution of RG. The three replicates are shown using three variants of the same color.

**Fig S5.**
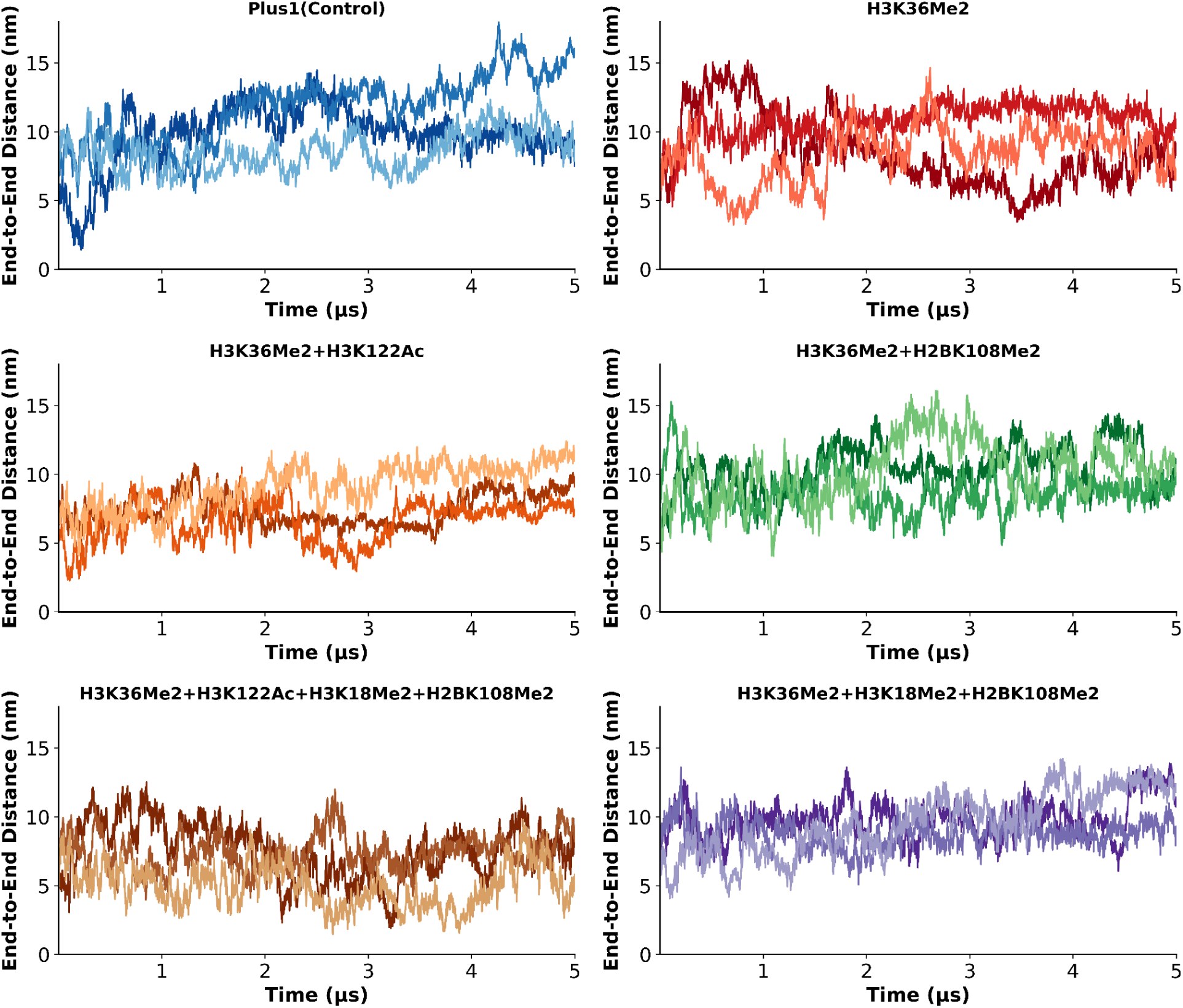
Depicts time evolution of End-to-end distance of DNA ends. The three replicates are shown using three variants of the same color.

**Fig S6.**
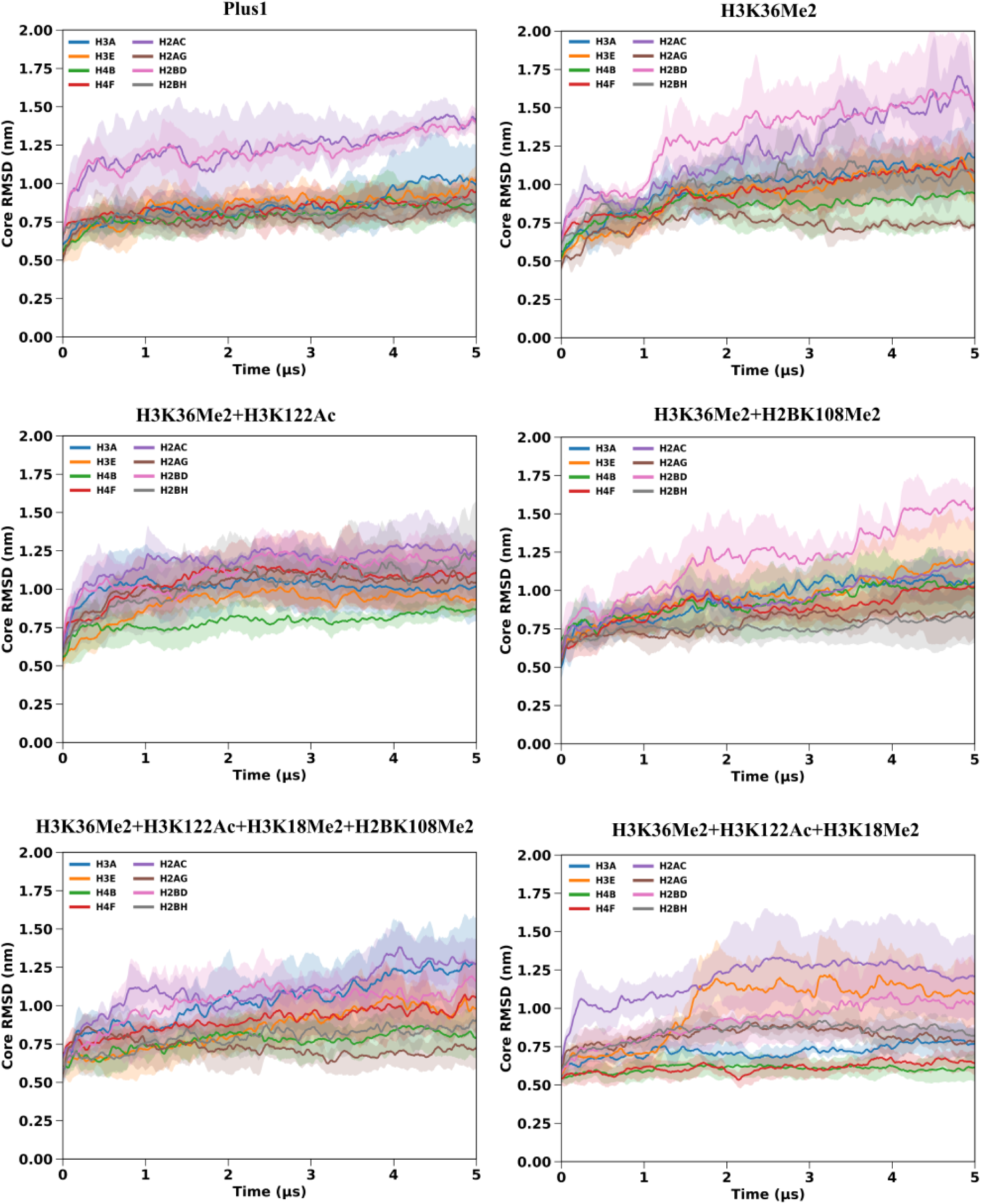
Depicts time evolution of RMSD of individual histone globular chains. The solid lines are the means and the shaded regions are standard deviations calculated using the three replicates.

**Fig S7.**
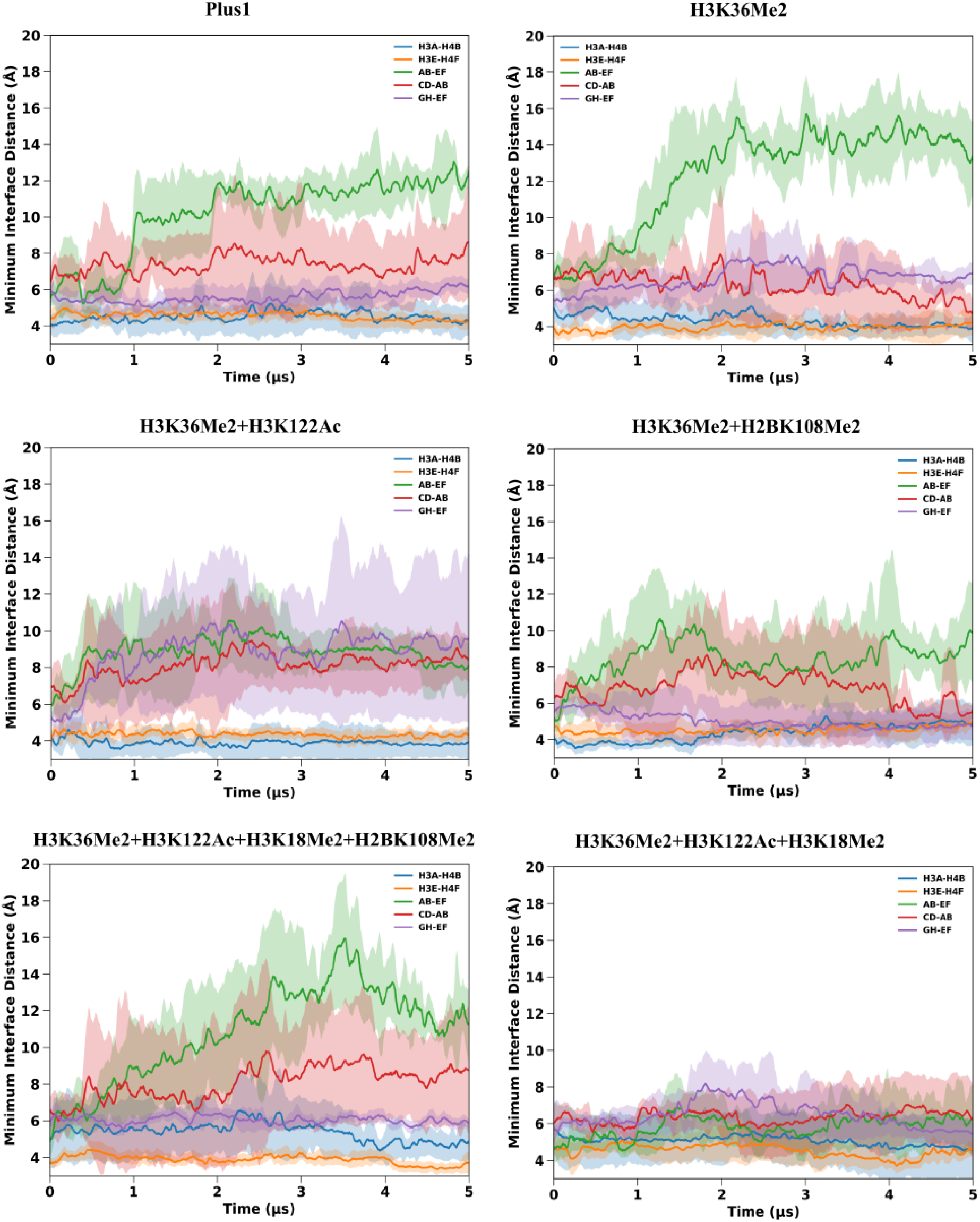
Depicts time evolution minimum interface distances between histone chains. The solid lines are the means and the shaded regions are standard deviations calculated using the three replicates.

**Fig S8.**
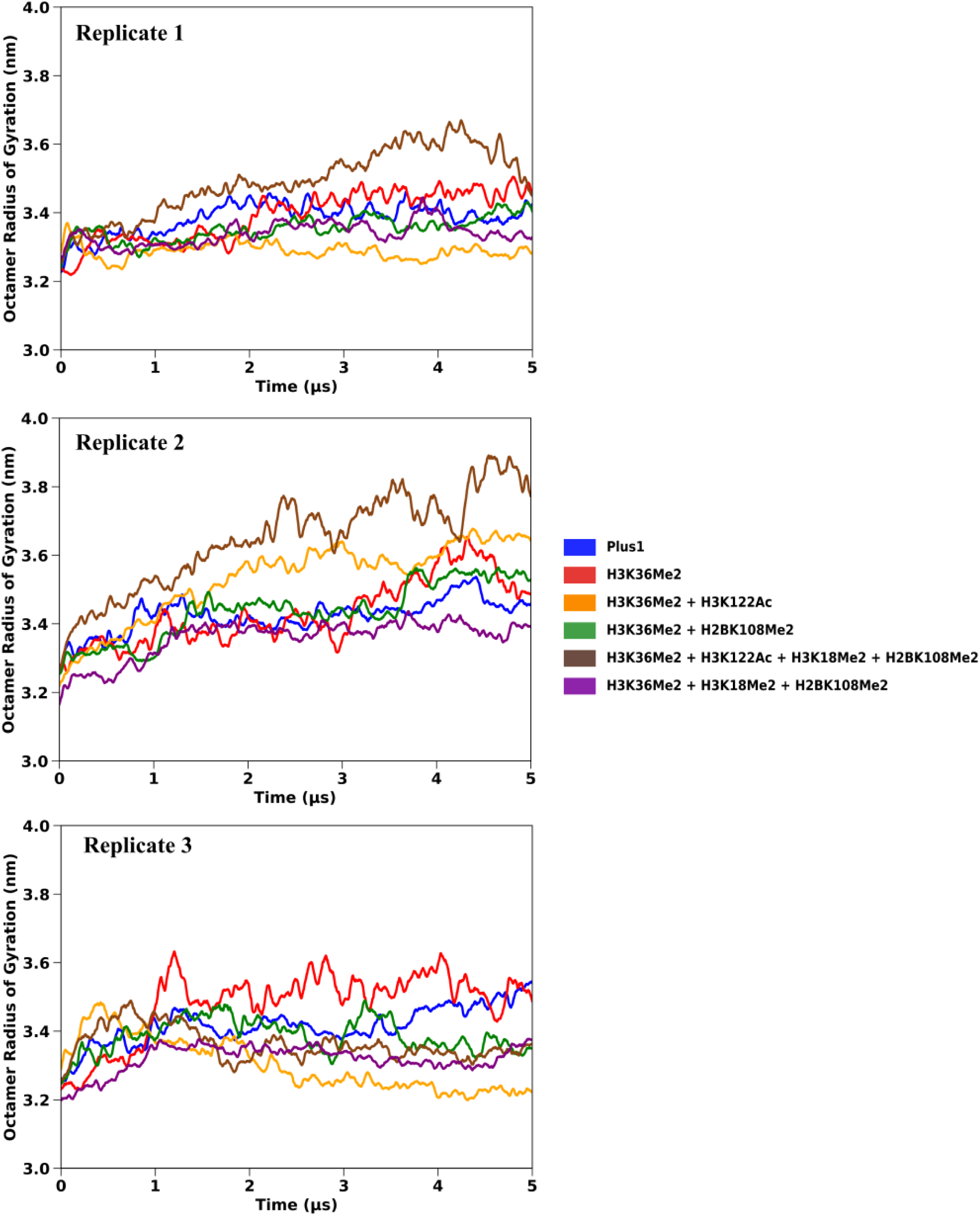
Depicts time evolution of RG of histone octamers.

**Fig S9.**
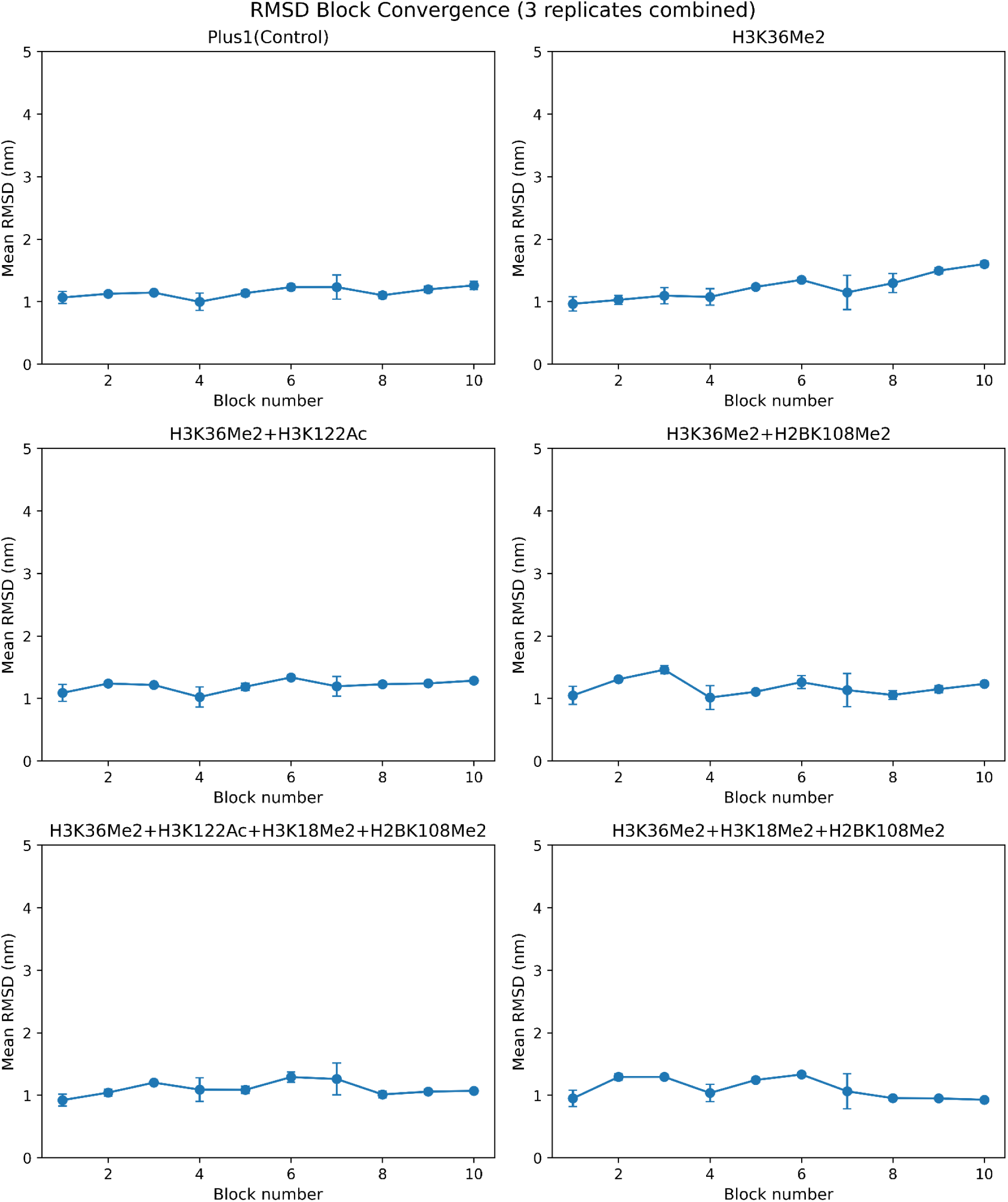
Block convergence analysis plots for RMSD of histone globules.

**Fig S10.**
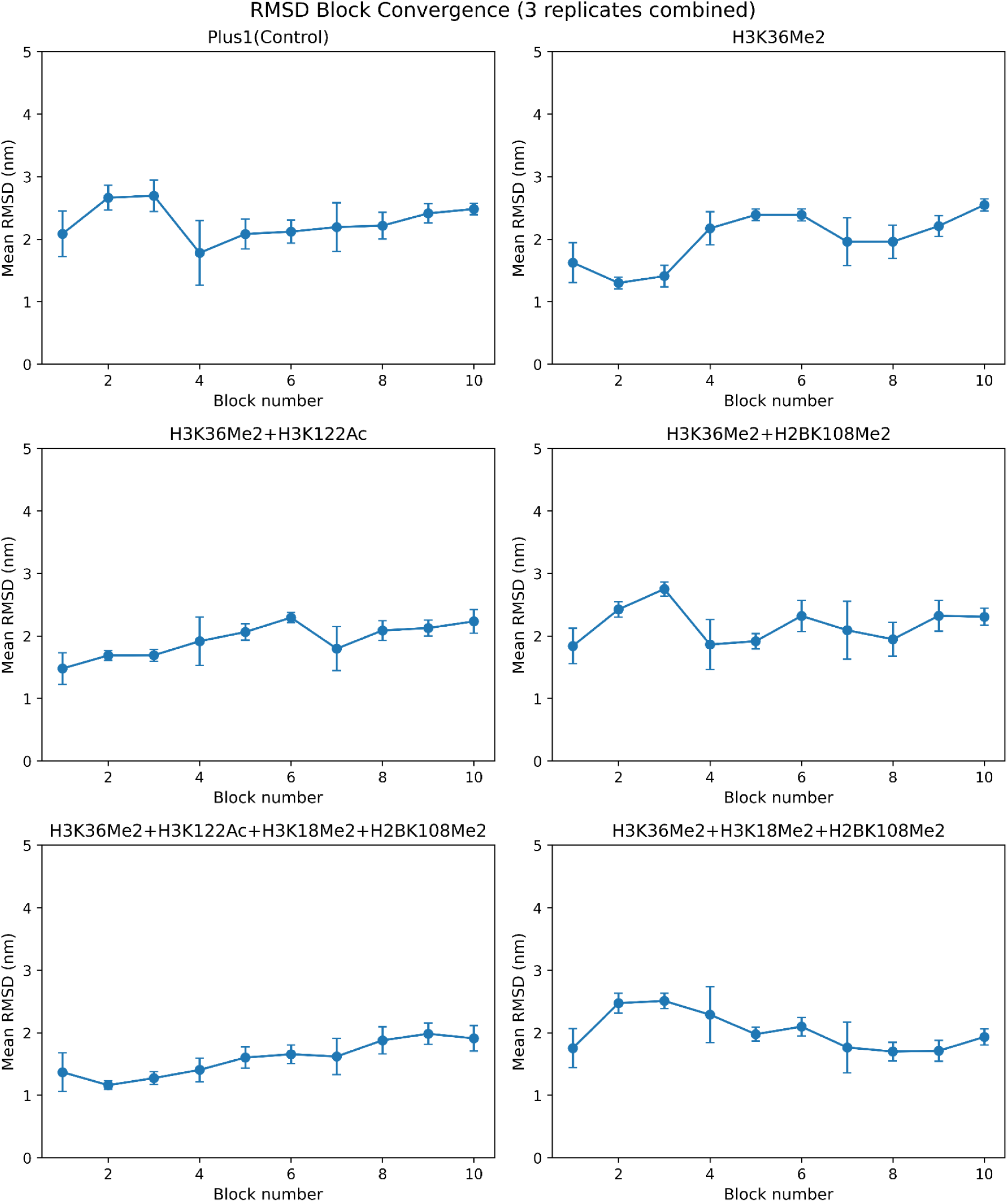
Block convergence analysis plots for RMSD of nucleosome DNA.

**Fig S11.**
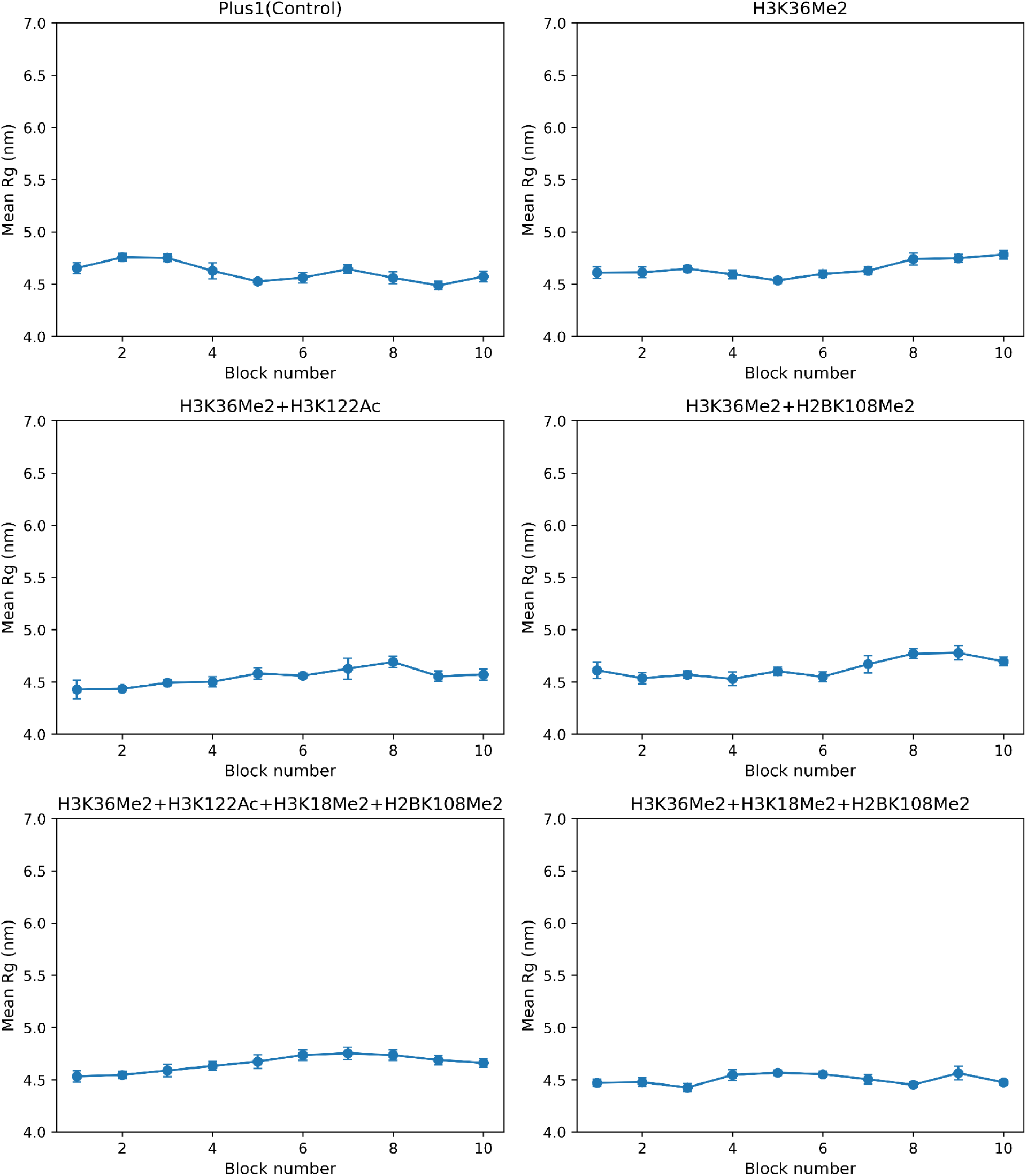
Block convergence analysis plots for RG of all systems.

**Fig S12.**
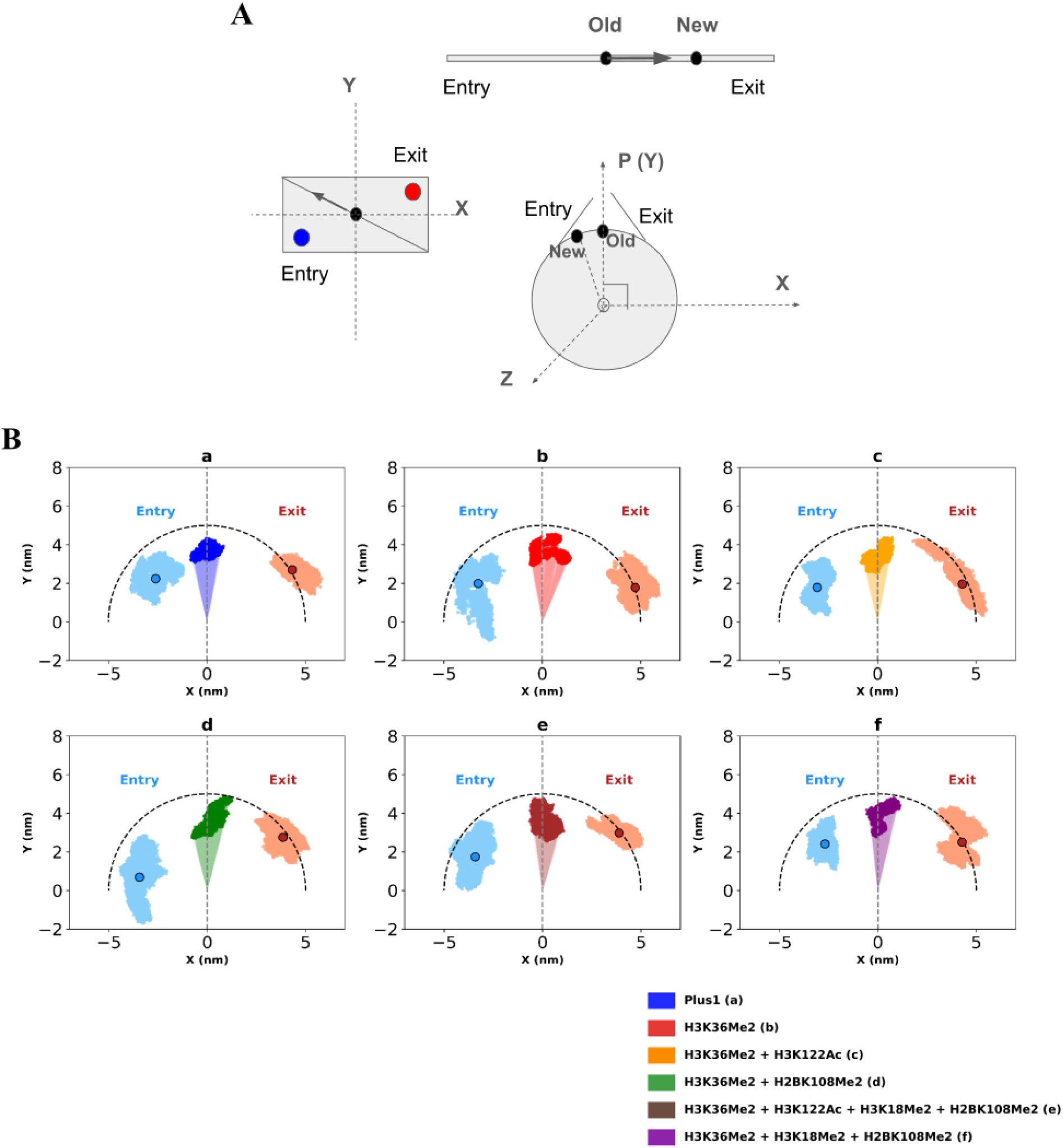
A) A schematic diagram of how the dyad sliding angles and dyad projection onto the X-Y plane is related to the nucleosome sliding direction. B) The dyad projection calculated based on the vectors considered in the work of Niina *et al.,* 2017.

**Fig S13.**
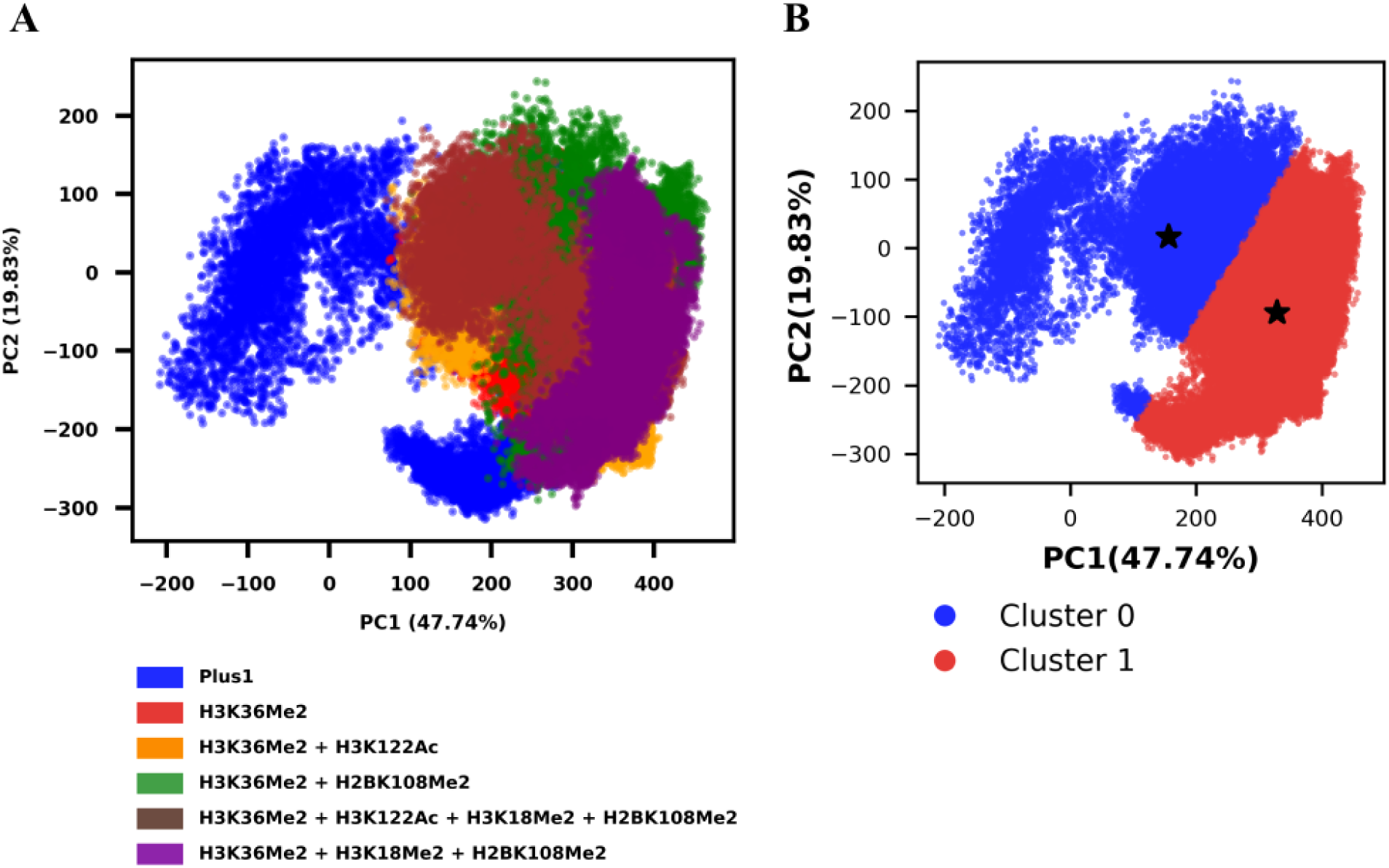
A) Principal component analysis (PCA) plot depicts DNA trajectory conformation of the nucleosome systems along the first two components PC1 and PC2 accounting for 47.74% and 19.83% of the total variance respectively. B) Depicts the two broad clusters into which all PCA clusters could be divided by k=2 means clustering. The two stars denote the representative “centroid” structures shown in Fig 4C of the main text.

**Fig S14.**
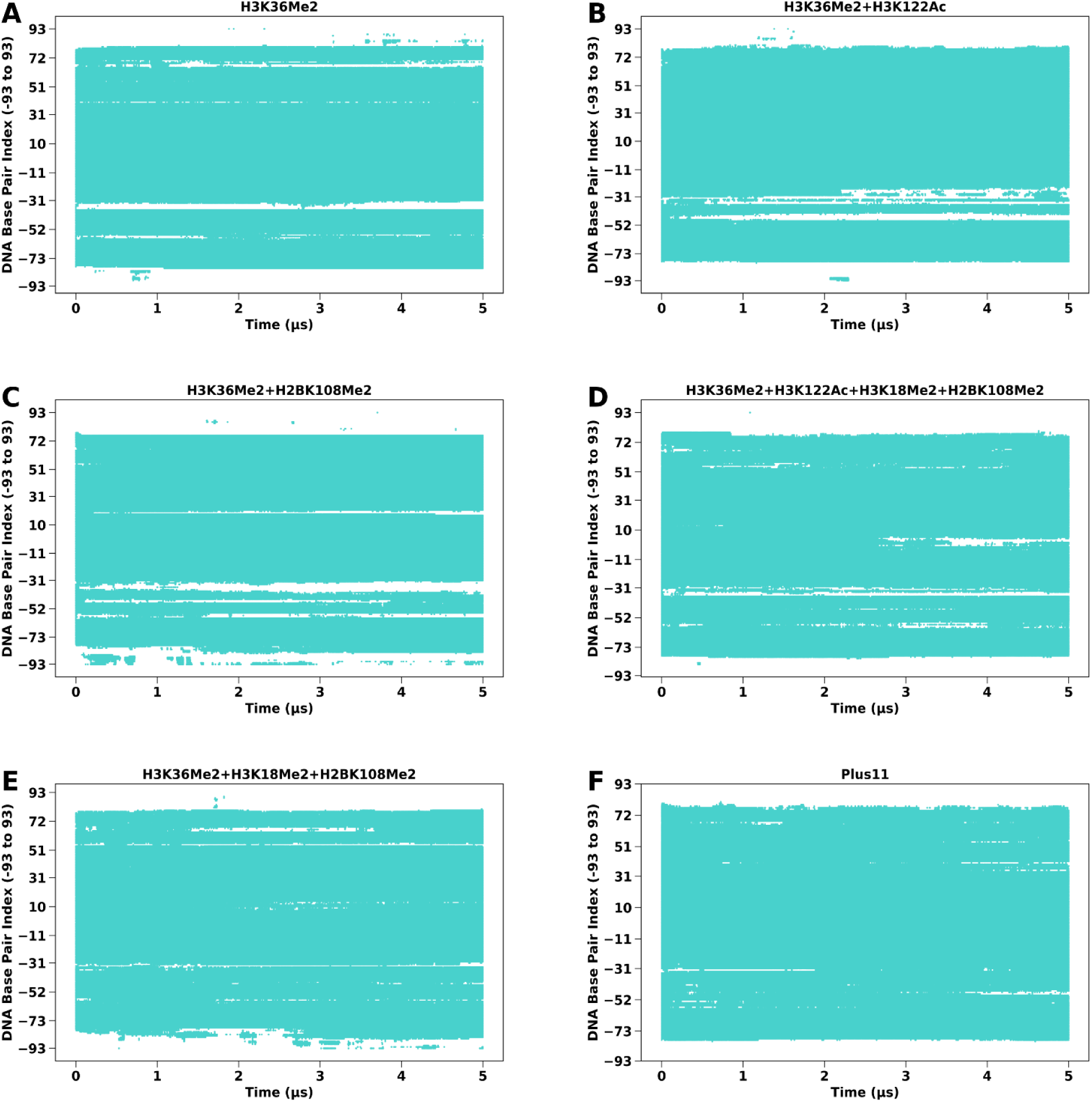
Binary contact maps of whole histone to DNA (contact if distance < 1 nm).

**Fig S15.**
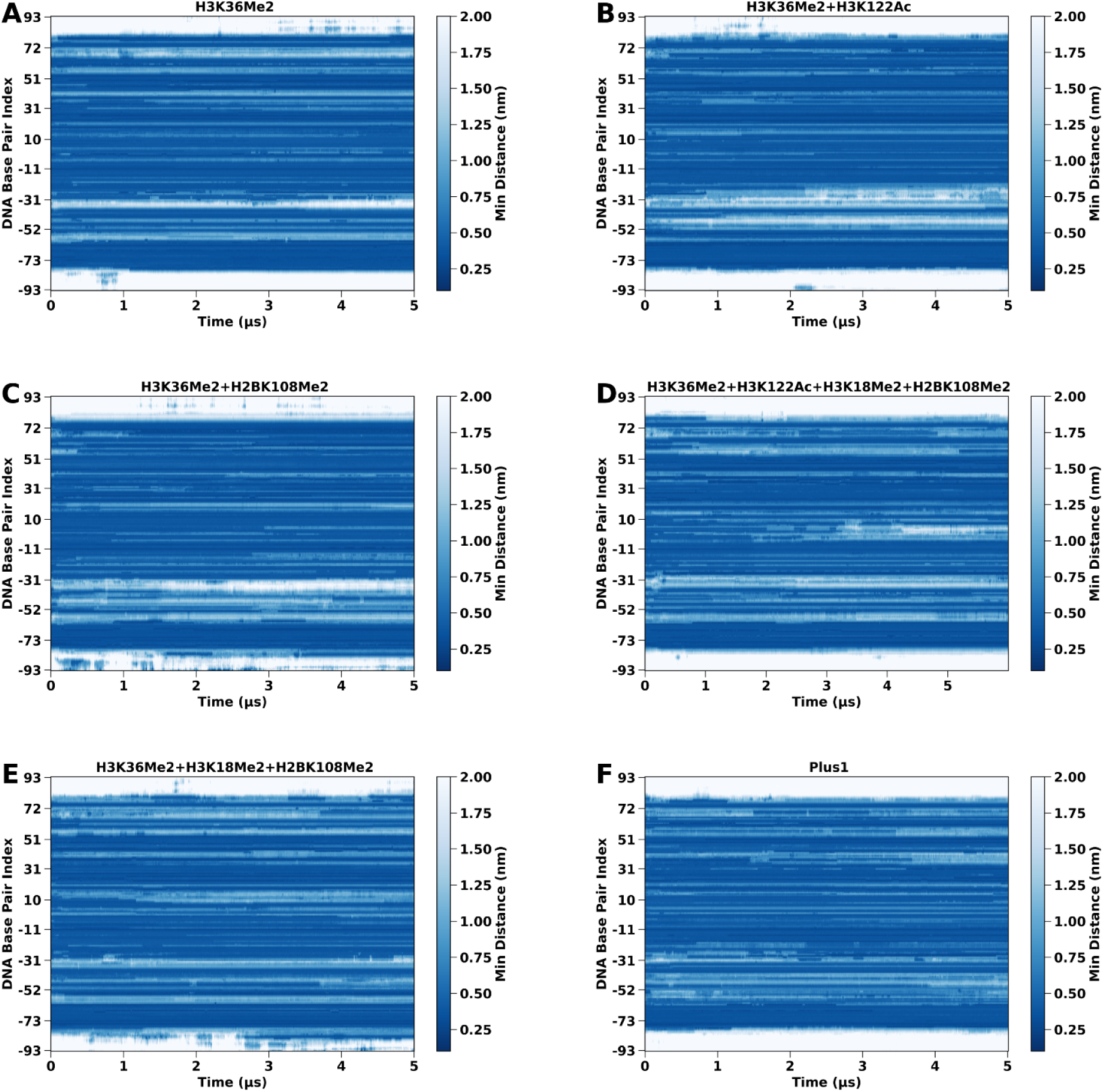
Contact heatmaps of whole histone to DNA.

**Fig S16.**
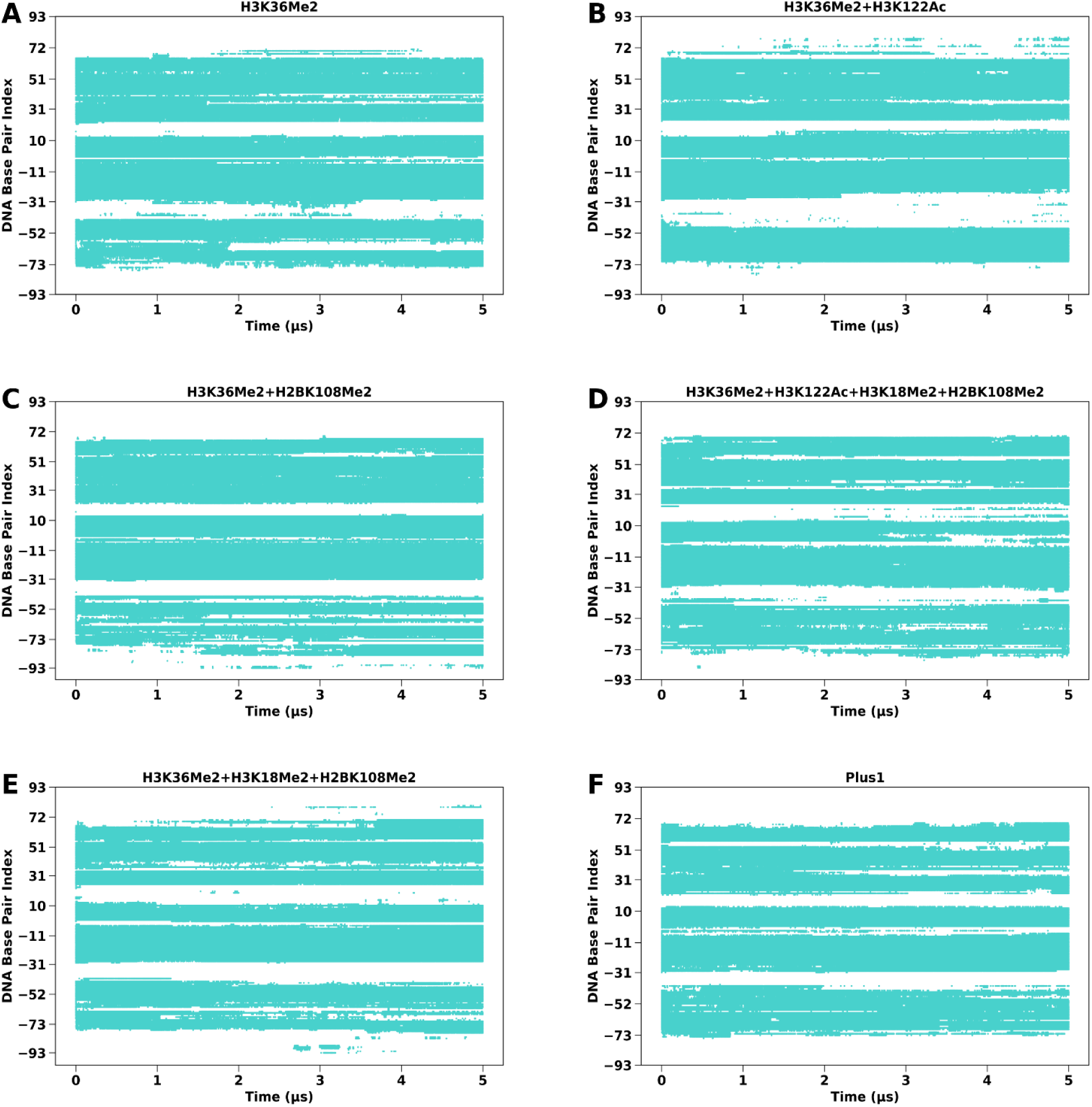
Binary contact maps of histone globular domain to DNA (contact if distance < 1 nm).

**Fig S17.**
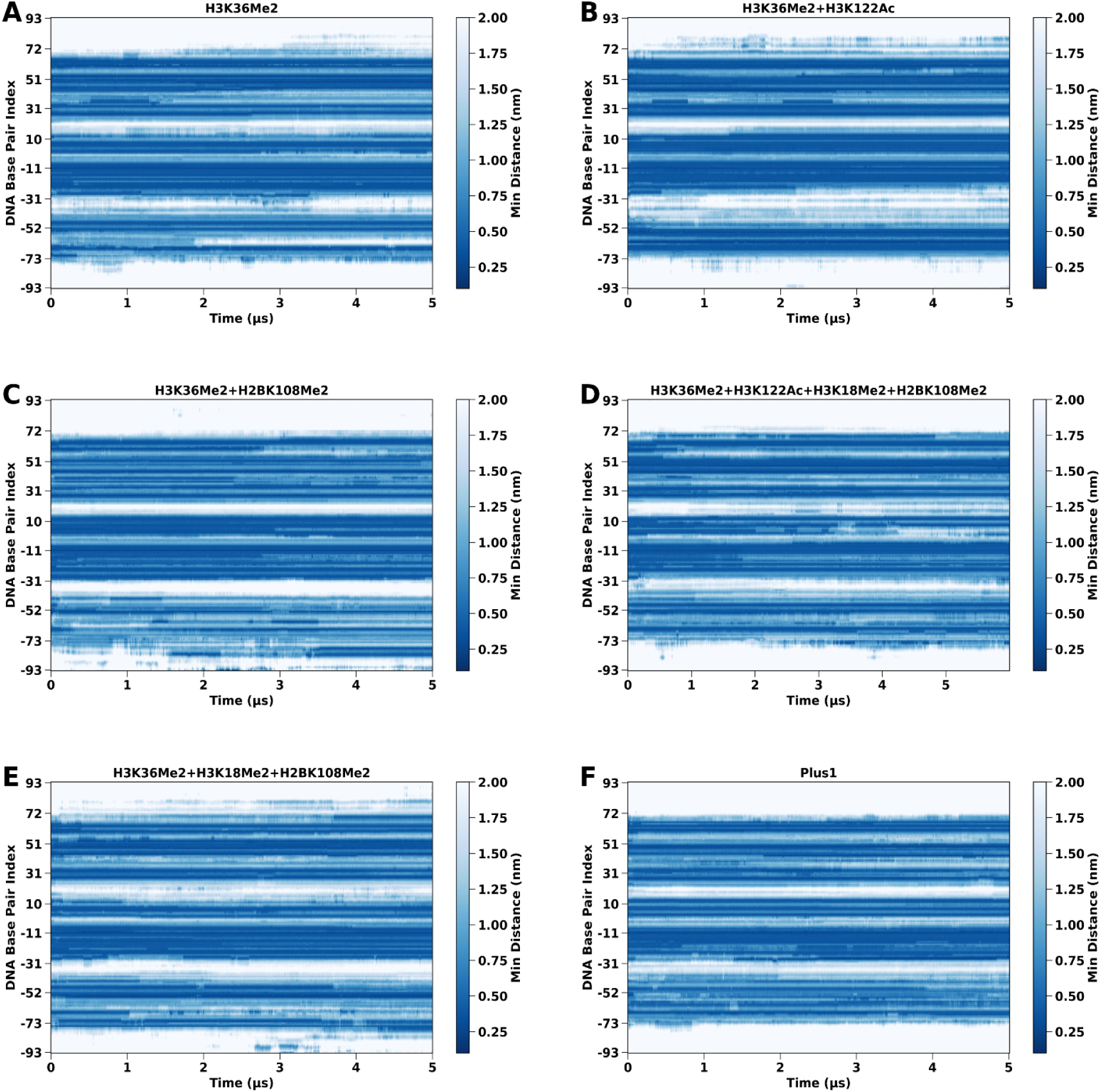
Contact heatmaps of histone globules to DNA.

**Fig S18.**
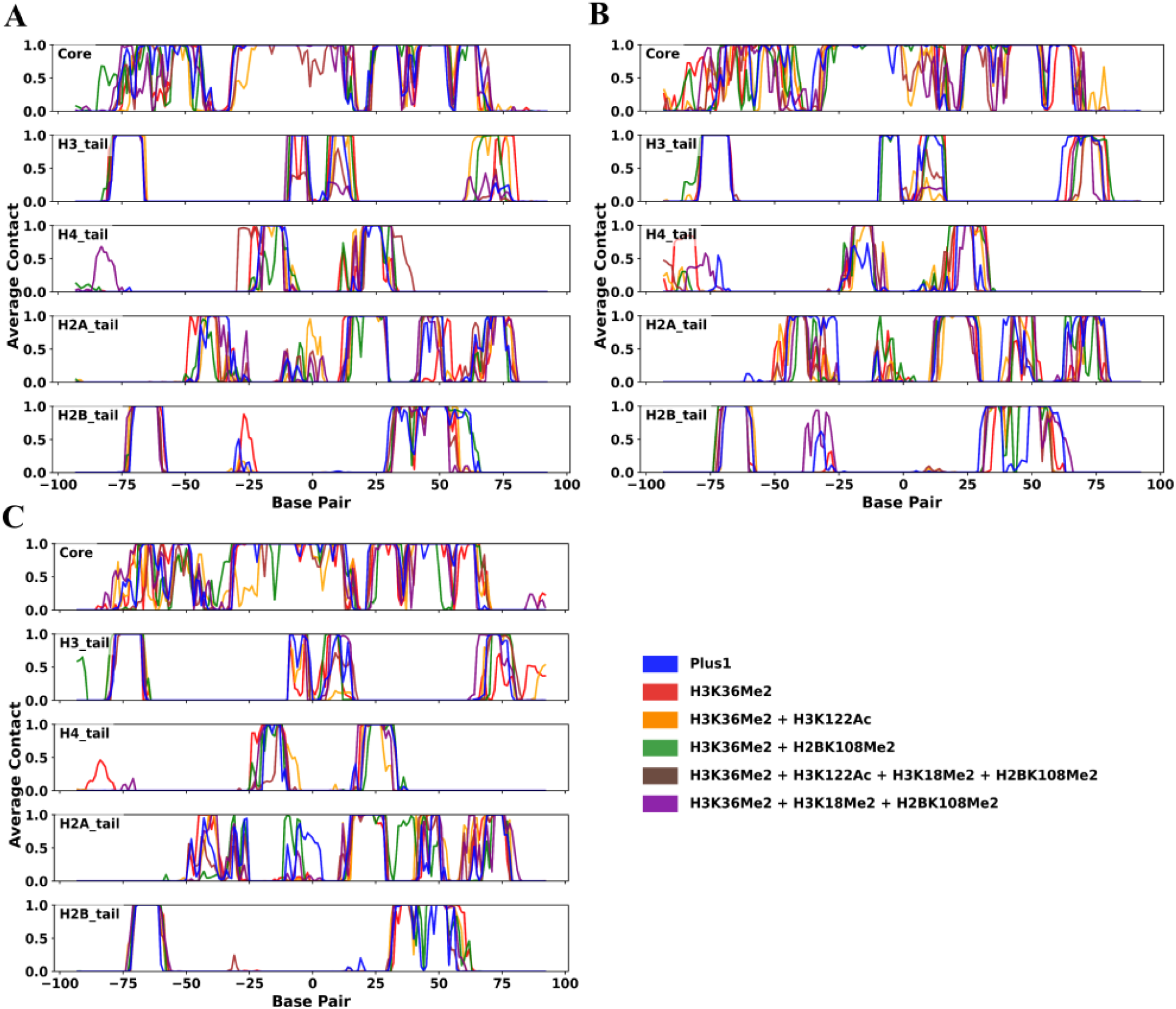
A-C) Average DNA contact profiles of histone core and tail regions across nucleosome systems. Each line represents the contact probability of the indicated system per DNA basepair, highlighting the difference of DNA-histone contacts at the DNA entry exit regions of the nucleosome.

**Fig S19.**
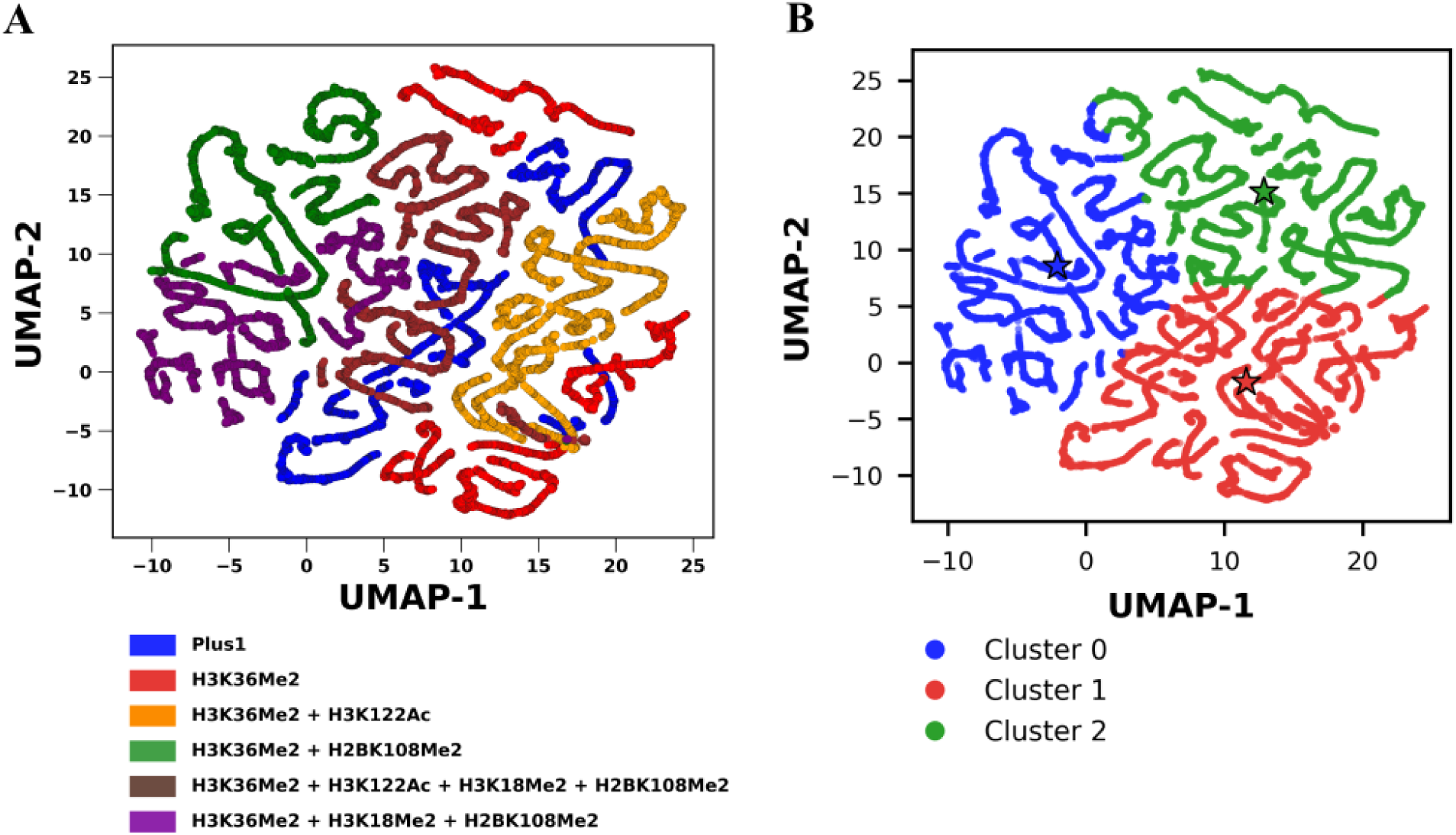
A) Uniform Manifold Approximation and Projection (UMAP) analysis based on the DNA-histone contact features, illustrating the clearly separated clusters into distinct conformational states. B) Shows the three clusters for UMAP with k-means (k=3) clustering. The three stars denote the representative “centroid” structures shown in Fig 5C of the main text.

